# Advancing single cell technology: iSCseq drives living subcellular transcriptomic profiling in osteoimmune diversity

**DOI:** 10.1101/2022.09.05.506360

**Authors:** Hiroyuki Okada, Yuta Terui, Masahide Seki, Asuka Terashima, Shoichiro Tani, Daisuke Motooka, Sanshiro Kanazawa, Masahiro Hosonuma, Junya Miyahara, Kenta Makabe, Yasunori Omata, Shoko Onodera, Fumiko Yano, Hiroshi Kajiya, Francesca Gori, Taku Saito, Koji Okabe, Yutaka Suzuki, Roland Baron, Ung-il Chung, Sakae Tanaka, Hironori Hojo

**Author notes:** Correspondence. (HO).

## Abstract

Single-cell RNA-seq (scRNA-seq) has clarified cellular heterogeneity within cell populations. However, scRNA-seq and spatial transcriptomics cannot capture the dynamic transcriptomic changes inside living cells. To decode subcellular gene expression, we developed intra-single cell sequencing (iSCseq), a novel approach that combines confocal imaging, repeatedly picking up cellular components inside living cells, and next-generation sequencing (intra single-cell RNA-seq; iSCseq). iSCseq illustrated the subcellular heterogeneity of gene expression. iSCseq revealed not only multiple differentiation stages embedded in the same cell, but also physical cytoskeletal connections, physiological activity of mitochondria, and intracellular calcium, as confirmed by transcriptomic evidence. Inclusive iSCseq with *in vivo* scRNA-seq datasets identified new osteoclast subsets in physiological and pathological bones. Network analysis with centrality provided insights into the connection between subcellular components, and clearly divided differentiation and fusion processes in multinucleation. The iSCseq approach has the potential to enhance cell biology at subcellular resolution and identify new therapeutic targets.

**Graphical abstract:** 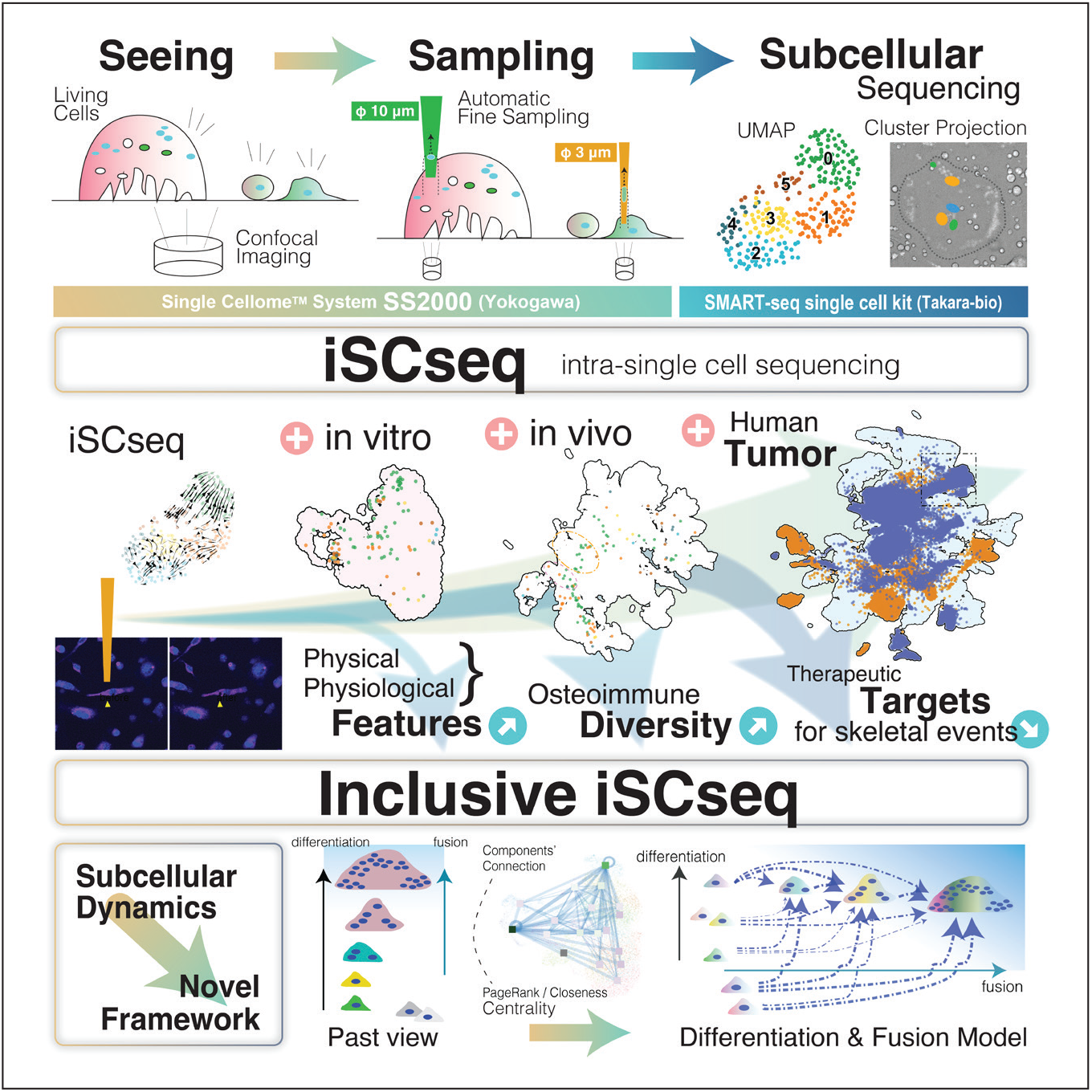

**In brief:** intra-single cell sequencing (iSCseq) enhances single-cell technology by combining live cell imaging, subcellular sampling from living cells and sequencing, offering deeper insights into cell functions and pathology at subcellular resolution through inclusive analysis with scRNA-seq and advanced centrality-focused network analysis.

**Highlights:**

- intra-single cell sequencing (iSCseq) clarifies subcellular heterogeneity
- iSCseq connects morphological and physiological features with transcriptome
- Inclusive iSCseq unveils osteoclast subsets in physiological and pathological bones
- Linkage at subcellular resolution reveals key players in characteristic fusion

## INTRODUCTION

Recent single-cell transcriptomic studies have clarified the cellular diversity within the cell population^1^. Spatial transcriptomics using fixed tissues connects transcriptomics with positional information on histological specimen^2,3^. However, these technologies cannot decode the dynamic localization of mRNA inside living cells. Live-seq challenged the recording of transcriptomes from living cells; however, it did not detect subcellular differences in gene expressions^4^. Moreover, it was difficult to elucidate the physiological and pathological implications of sequencing at subcellular resolution.

During the bone remodeling process, hematopoietic and mesenchymal cells are coordinated to maintain bone mineral density (BMD)^5^. Fails in cooperative relationships lead to skeletal-related events such as geriatric, inflammatory, and pathological fractures, which are global challenges in the aged society^6,7^. Although bone resorption inhibitors increase BMD and reduce the fracture risk, their use is often prohibited in patients with renal failure. There remains global controversy regarding the withdrawal of anti-bone resorptive reagents, sometimes causing rapid bone loss^8^. Further investigation of the mechanism of bone resorption is necessary to address the unmet clinical needs.

Osteoclasts (OCs), which have characteristic large portion and often contain >10 nuclei, are hematopoietic stem cell-derived bone-resorbing cells induced by macrophage colony stimulating factor (M-CSF; *CSF1*) and receptor activator of nuclear factor kappa B ligand (RANKL; *TNFSF11*)^9^. Nuclear factor of activated T-cells (NFATc1) is a master regulator of OC differentiation^10^. OC is highly polarized, with a specific conformation of its cytoskeleton in an actin ring that seals off an extracellular bone-resorbing compartment^11^. This specific cellular organization in the presence of multinucleation raises the question of whether nuclei residing in the same cytoplasm are similarly regulated or heterogeneous, expressing a different set of genes and/or different levels of physiological activity. Essential signaling cascades inside cells, such as mitochondrial metabolism and fine-tuning of calcium concentration inside OC^12-14^, have not been detected, but are predicted. In addition, the difference between *in vitro* cultured OC and *in vivo* OC remains unclear because of the difficulty in sampling from the bone. Moreover, the origin of OCs in the hematopoietic lineage has not been quantitatively examined.

To clarify the heterogeneity of OCs, scRNA-seq was performed for *in vitro*-cultured OCs^15-17^. However, conventional single-cell techniques theoretically do not allow for the separation of OCs into single-cell suspensions because of their large size and fragility. Although there is no cell-by-cell transcriptomic evidence for giant cells, differentiation and fusion have been discussed. Therefore, new platforms must be developed to reveal the diversity of OCs.

This study aimed to establish a new technology called intra single cell sequencing (iSCseq) to examine subcellular transcriptomic heterogeneity in living OCs and small cells. Matching the data points obtained by iSCseq with conventional scRNA-seq datasets clarified the physiological and pathological mechanisms of bone resorption in humans.

## RESULTS

### The basics of iSCseq (intra-single cell sequencing)

Intra single cell sequencing (iSCseq) is a new approach that combines confocal live cell imaging, intracellular sampling with a 3-μm or 10-μm probe, and next-generation sequencing from ultra-pico input of RNA using commercially available kits. iSCseq enabled us to capture the transcriptomes of cellular and subcellular components, including individual nuclei.

Mature OCs are large cells with diameters of > 100 μm *in vitro*. Single Cellome™ System SS2000 (Yokogawa, Japan) enabled the sampling of not only a single cell, but also subcellular components inside living OCs under visual observation. Using this approach, we sampled single nuclei from the peripheral zone of the OC (Figure 1A, Movie S1) or one nucleus from the aggregation of four nuclei (Figure 1B, Movie S2). Fine-tuning of the probe tip position and negative pressure also enabled multiple samplings from ordinary small single-nucleated cells (Figure 1C). iSCseq sampled a maximum of 11 batches from one OC and 15 batches from one foreign body giant cell (FBGC, Figure S1D). FBGCs induced by IL-4 are derived from the same monocyte lineage as OC; however, FBGCs do not resorb bone^18^.

**Figure 1.**
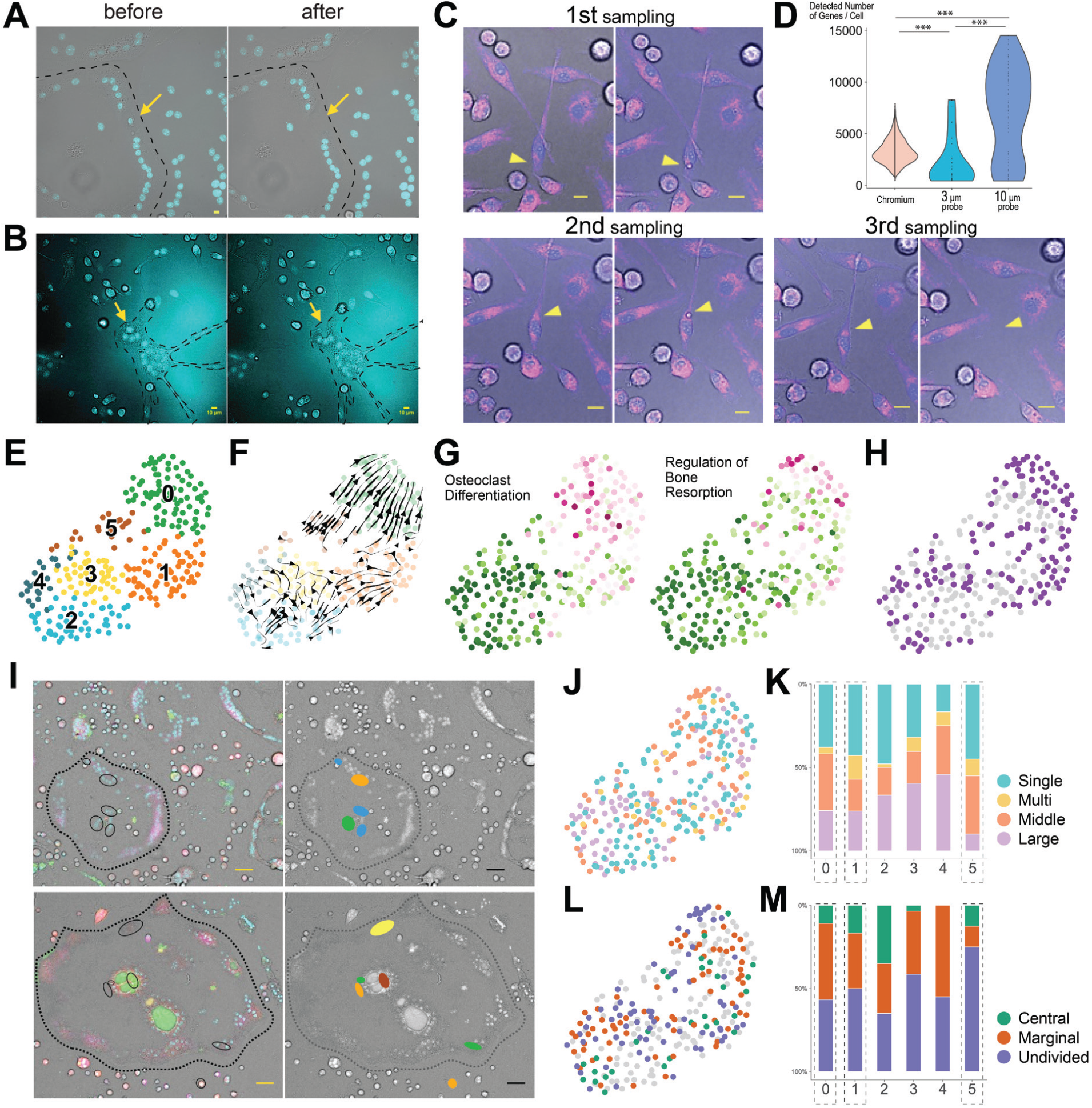
intra-single cell sequencing (iSCseq) for cellular and subcellular components. (A-C) Representative cases of sampling each nucleus or subcellular component from a single cell. Blue: nuclei stained with Hoechst 33342. Bar = 10μm. (A, B) The Area enclosed by the black dotted line represents a single osteoclast. (A) Nucleus from the peripheral zone of a large osteoclast (OC). (B) A nucleus from the aggregation of four nuclei in an OC. (C) 1st and 2nd sampling of each subcellular component. 3rd sampling of a whole cell with a nucleus. Purple: mitochondria stained with MitoBright LT Deep Red. (D) Violin plot of genes detected by the sampling and sequencing methods. Left to right, fluid-based chromium (10x genomics), SMART-seq for samples picked by 3 μm probe, and by 10 μm probe. ***p<0.001, One-way ANOVA and Tukey post hoc test. (E) UMAP consisted of 280 cellular and subcellular components. (F) Streamline presentation based on the spliced/unspliced ratio of RNA using scVelo. (G) Gene ontology terms ‘Osteoclast Differentiation’ and ‘Regulation of Bone resorption’ analyzed using decoupleR and AUCell. (H) Automatic cell type annotation using CellAssign. Purple: Osteoclast. (I) Geometrical information of the targeted mature OCs (left). Blue: nuclei stained with Hoechst 33342; Green: Calcium stained with Fluo-4 NW Calcium Assay Kit; Red: acidic environment stained with pHrodo™ Red AM Intracellular pH Indicator; Purple: Tubulin stained with SiR-tubulin kit. Projection of the iSCseq cluster (right). Bar = 50 μm. (J, K) Nuclear Rank: bin, number of nuclei in the origin of cells. Large, 20 or more nuclei; Middle, 5 to 19 nuclei; Multi, 2 to 4 nuclei; Single, 1. (J) Feature plot of Nuclear Rank. (K) Cumulative bar plot of Nuclear Rank in each cluster. OC-enriched clusters are surrounded by dotted rectangles. (L, M) Positional information, Central or Marginal or Undivided in OC. (L) Feature plot of the positional information. (M) Cumulative bar plot of positional information for each cluster. OC-enriched clusters are surrounded by dotted rectangles.

Next, we connected fine samples of cellular or subcellular components confirmed by multicolored confocal images with transcriptomic library construction using SMART-seq single cell PLUS kit (Takara Bio, Japan). The sampled cellular or subcellular components were harvested in a microtube containing RNA lysis buffer with RNase inhibitors, centrifuged, and frozen promptly to minimize RNA deterioration. The number of genes detected with a 10-μm probe were more than twice those detected with conventional fluid-based chromium (10x genomics, U.S.) (Figure 1D).

### iSCseq connects gene expression with morphology

Gene expression from 280 of 285 samples (98.2 %) was divided into six clusters on UMAP^19^ (Figure 1E) using Python-based deep learning scvi-tools^20-22^, considering the percentages of mitochondrial reads (%mt), ribosomal reads, and cell cycle phase defined by the Seurat team^23^ (Figure S2A-C). The differentiation pathway was examined based on the RNA-spliced/unspliced ratio using scVelo^24^ (Figure 1F). Gene ontology (GO) analysis using decoupleR^25^ (Figure 1G) and automatic cell type annotation using CellAssign^26^ (Figure 1H) illustrated that cellular components in clusters 0, 1, and 5 showed an OC phenotype. Cluster 0 was regarded as the terminal cluster in OC differentiation based on the cell-type annotation (Figure 1H, S1E and S1F) and OC-related gene expression and GO terms (Figure S1G and S1H). The clustering results were projected onto the captured images, which provided morphological annotations (Figure 1I).

Next, we reviewed the number of nuclei in the original OC using binned Nuclear Rank. Cluster 0 consisted of fewer subcellular components from large OCs with 20 or more nuclei, which were morphologically called OCs, than the average proportion (Figure 1J and 1K). Geographical positions were dichotomized into central or marginal nuclei. Cluster 0 contained more central nuclei than clusters 1 and 5 (Figure 1L and 1M).

These results demonstrated the subcellular heterogeneity in the same mature OC. Some single-nucleated small cells, which were believed not to be OCs in the classical morphological definition, had already differentiated into the final OC cluster.

### Inclusive *in vitro* iSCseq combined with conventional scRNA-seq

To validate our iSCseq approach, we performed an integrative analysis with two previously published scRNA-seq studies on *in vitro* OC differentiation (inclusive *in vitro* iSCseq), which were generated using a traditional microfluid-based technique^15,17^. A total of 12,840 cellular or subcellular components, including 116 iSCseq results, with less than 25 %mt (Figure S2A) and more than 400 genes, were illustrated on UMAP (Figure 2A). Although cytokine treatment options were considered in the batch effect correction, cells in clusters 4 and 10 and almost all cells in cluster 17 appeared only under RANKL stimulation (Figure 2D and S2C). In inclusive iSCseq, the differentiation pathway was directed upward on the right island, and cluster 10 was regarded as the terminal cluster in OC differentiation (Figure 2B, 2C, 2E, S2D and S2F). The iSCseq spots rendered on inclusive *in vitro* UMAP depicted the differentiation stage of each cellular component (Figure 2F). The terminal OC cluster was selected more strictly by inclusive iSCseq than by iSCseq analysis alone (Figure 2G). Nuclear rank analysis showed that OC clusters in inclusive *in vitro* iSCseq did not contain a higher than average proportion of fragments from large OCs, which were believed to be mature OCs (Figure 2H and 2I). Interestingly, OC clusters included ordinary single-nucleated cells (Figure 2J). Nucleus positional analysis showed that cluster 10 contained fewer central nuclei than cluster 4 (Figure 2K and 2L). In a representative OC, the cluster number differed according to the central or marginal position (Figure 2M).

**Figure 2.**
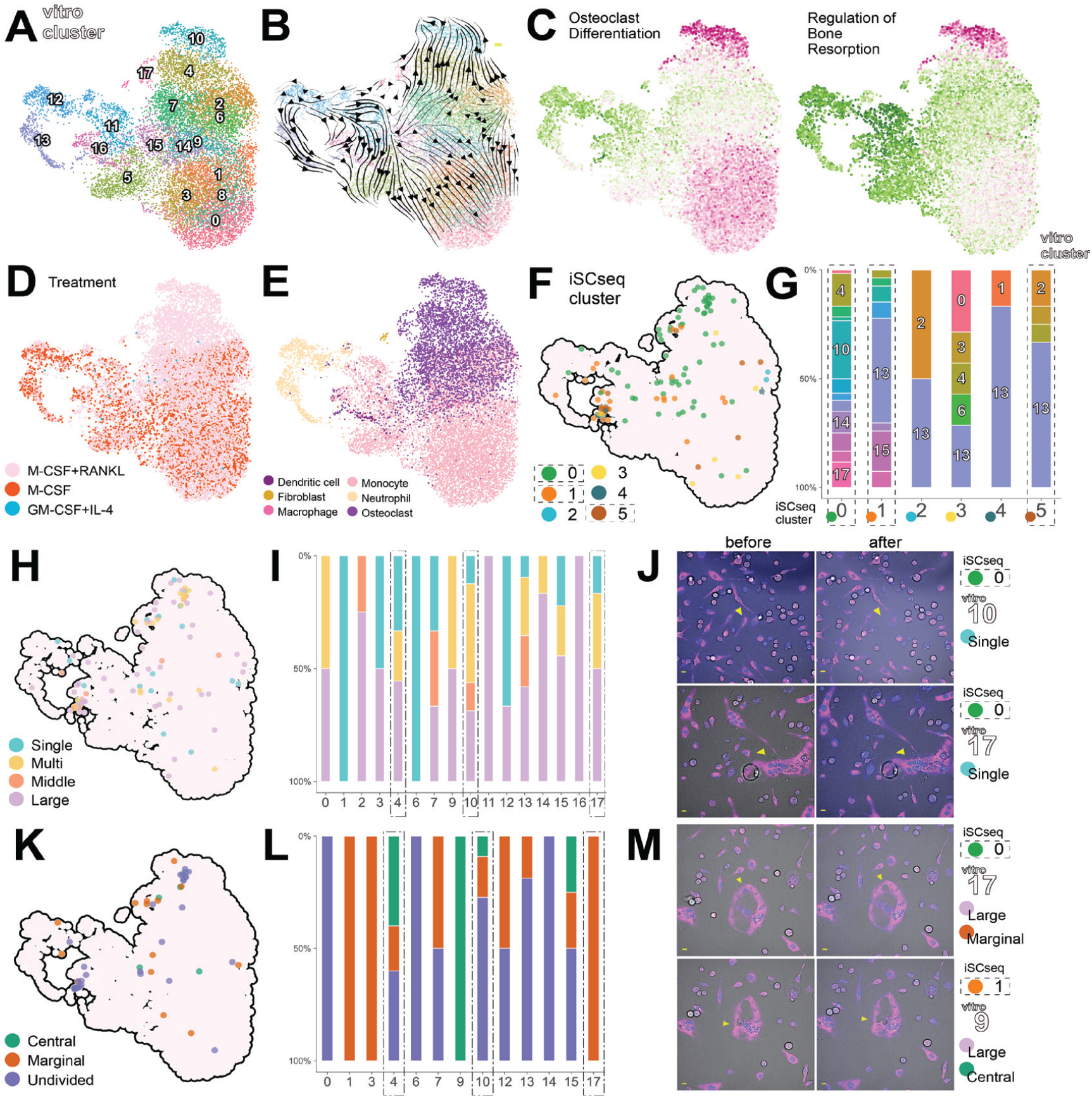
Inclusive *in vitro* iSCseq combined with conventional scRNA-seq. (A) Inclusive *in vitro* iSCseq UMAP comprising iSCseq and scRNA-seq OC datasets. (B) Streamline presentation based on RNA velocity using scVelo. (C) Gene ontology ‘Osteoclast Differentiation’ and ‘Regulation of Bone resorption’ analyzed using decoupleR and AUCell. (D) Feature plot of the inductive treatment of cytokines. (E) Automatic cell type annotation using CellAssign. (F) iSCseq spots on inclusive *in vitro* UMAP. Colors corresponding to the clusters obtained by iSCseq are shown in Figure 1. (G) Cumulative bar plot of the inclusive *in vitro* clusters for each iSCseq cluster. OC clusters in iSCseq are indicated by dashed rectangles. (H, I) Nuclear Rank: bin, number of nuclei in the origin of the cell. Large, 20 or more nuclei; Middle, 5 to 19 nuclei; Multi, 2 to 4 nuclei; Single, 1. (H) Feature plot of Nuclear Rank. (I) Cumulative bar plot of Nuclear Rank in each cluster. OC clusters of inclusive *in vitro* iSCseq are surrounded by dotted-dashed rectangles. (J) Representative cases of single-nucleated cells assigned to OC clusters in *in vitro* iSCseq. (K, L) Positional information of samples, Central or Marginal or Undivided in OC. (K) Feature plot of positional information. (L) Cumulative bar plot of positional information for each cluster. OC clusters of inclusive *in vitro* iSCseq are surrounded by dotted-dashed rectangles. (M) Representative sampling cases of a nucleus from a multinucleated OC. Blue, Nuclei stained with Hoechst 33342; Purple, MitoBright LT Deep Red. Bar = 10 μm.

Inclusive iSCseq depicted the traits of subcellular components at a higher resolution than iSCseq alone (Figure S2E). Even with more strict definition, some small single-nucleated cells were categorized into terminal OCs.

### Interventional response evaluated by iSCseq

iSCseq precisely controls the tip of the probe at the xy and z positions. In addition, we were able to manipulate the duration and negative pressure sample by sample. Although the sampling procedure was tuned and equalized, multiple nuclei were occasionally collected using a probe (Figure 3A-C). We hypothesized that the sampling reaction could reflect the tightness of the cytoskeleton among the nuclei and be evaluated in terms of gene expression.

**Figure 3.**
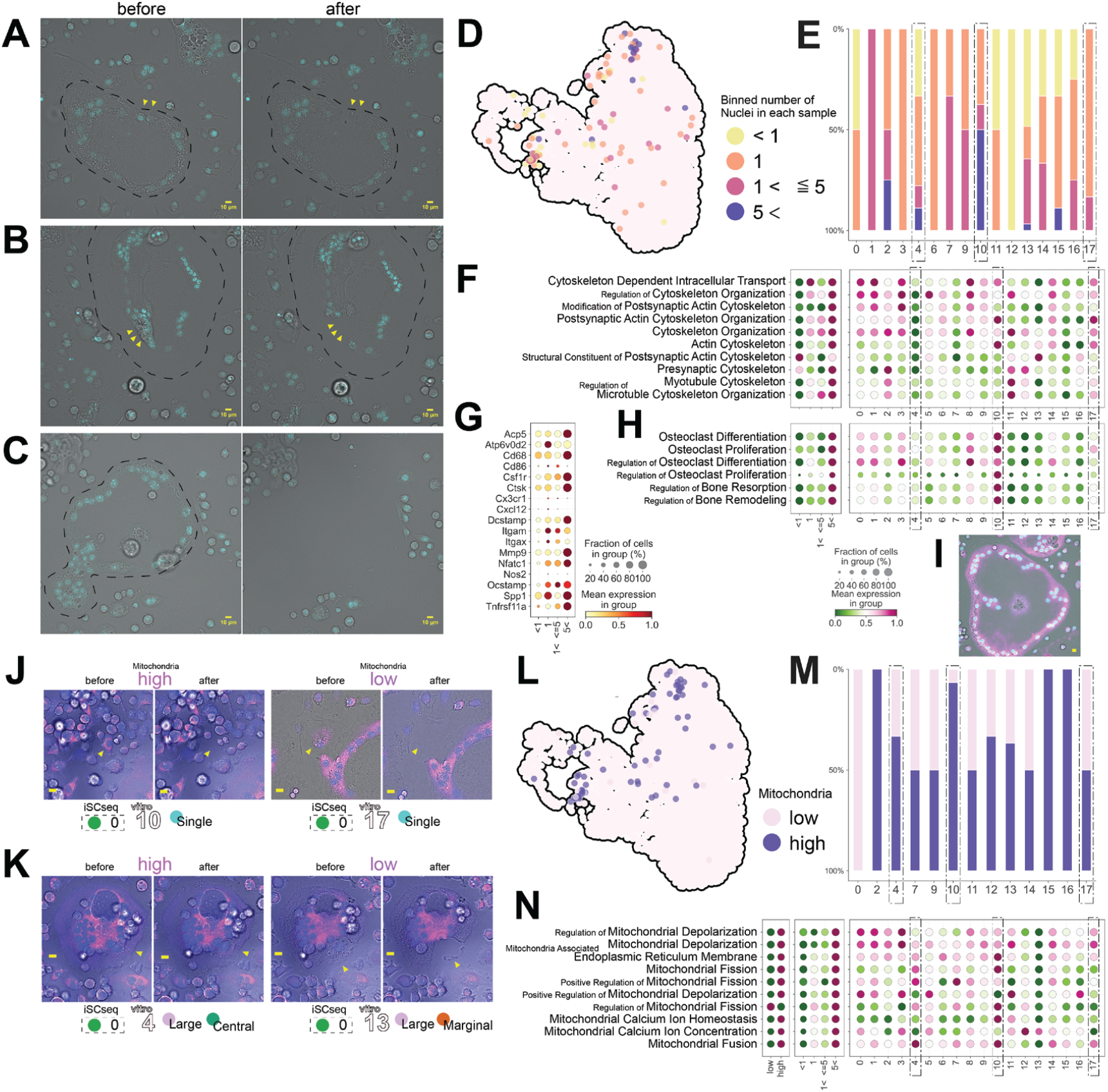
Morpho-physiological analysis of physical responses and of mitochondria with transcriptome. (A-C) Representative nuclei sampling cases from multinucleated cells. Blue: nuclei stained with Hoechst 33342. The area enclosed by the black dotted line represents one osteoclast (OC). Bar = 10 μm. (A, B) Yellow arrowheads indicate the sampling objects. (A) Two nuclei in the marginal zone. (B) Twelve nuclei from nuclei aggregations. (C) Sampling a whole large osteoclast. (D, E) Binned number of nuclei in each sample. Feature plot (D) and cumulative bar plot for inclusive *in vitro* cluster shown on Figure 2A (E). OC clusters are surrounded by dotted dashed rectangles. (F) Dot plot of the top 10 gene ontology (GO) terms including cytoskeleton by decoupleR and AUCell by binned number of nuclei in each sample (left) and by inclusive *in vitro* iSCseq cluster (right). OC clusters are surrounded by dotted dashed rectangles. (G) Dot plot of OC-related gene expression by the binned number of nuclei. (H) Dot plot of OC-related GO terms by binned number of nuclei in each sample (left) and by inclusive *in vitro* iSCseq cluster (right). (I) Representative image of a large OC. Blue, Nuclei stained with Hoechst 33342; Purple, Tubulin stained with SiR-tublin kit. Bar = 10 μm. (J, K) Representative sampling cases of single-nucleated cells (J) and multinucleated OC (K). Blue, Nuclei stained with Hoechst 33342; Purple, MitoBright LT Deep Red. Bar = 10 μm. (L, M) High- or low-dichotomized information of mitochondrial intensity from sampling images. Feature plot (L) and cumulative bar plot by inclusive *in vitro* iSCseq cluster (M). OC clusters are surrounded by dotted dashed rectangles. (N) Dot plot of the top 10 gene ontology terms including mitochondria by decoupleR and AUCell by high or low mitochondrial intensity from images (left), by inclusive *in vitro* iSCseq cluster (center), and by binned number of nuclei in each sample (right). OC clusters are surrounded by dotted dashed rectangles.

Analyses of the binned number of nuclei in each OC sample showed that final cluster 10 contained the most components with more than five nuclei (Figure 3D and 3E) and exhibited upregulation of cytoskeleton-related terms and OC phenotype (Figure 3F and 3H). Conversely, OC components with more than five nuclei showed a stronger OC phenotype and upregulation of cytoskeleton-related terms (Figure 3F, 3G and 3H).

Gene ontology analysis explained the physical reactions during the interventional sampling well. iSCseq provided transcriptomic evidence for living cell motion. In addition, cluster 10 contained nuclei-abundant ingredients. Considering figure 2H and 2I, this indicated that the terminal cluster contained a variety of components, ranging from mononuclear to multinuclear.

### Inspective morpho-physiological analysis with gene expression

It is difficult to evaluate mitochondrial genes using conventional scRNA-seq. Seurat used %mt as a normalization factor ^23,27,28^, which means that mitochondrial gene expression was assumed to be roughly equivalent. iSCseq can add transcriptome evidence to morphological and physiological findings based on live-cell imaging. We examined the relationship between mitochondrial gene expression in each fraction and dichotomized mitochondrial density based on fluorescent images obtained by iSCseq experiments.

Figure 3J and 3K were representative small and multinucleated cells whose mitochondria were stained purple, with the information of the inclusive *in vitro* iSCseq cluster. We can provide information on mitochondrial fluorescence density on inclusive *in vitro* iSCseq UMAP (Figure 3L). Terminal cluster 10 was rich in components with a high mitochondrial intensity (Figure 3M). Mitochondrial dichotomization, based on the images, was confirmed using GO terms (Figure 3N, left). Fractions containing more than five nuclei and the terminal cluster exhibited upregulation of mitochondrial-related terms (Figure 3N, center and right). Cellular calcium concentration was evaluated using the same scheme as that used for mitochondrial evaluation (Figure S3).

Evaluating the distribution of cellular organs or small molecules inside living cells is challenging. Live cell imaging using antibodies is strongly affected by experimental factors including antibody affinity. The iSCseq approach proved that gene expressions confirmed the physiological activities based on live cell images. The iSCseq results suggested an unequal distribution of intracellular organs even in small cells.

### Inclusive *in vivo* iSCseq with single cell bone atlas

iSCseq based on live-cell imaging has linked reality to gene expression using anonymous conventional single-cell RNA-seq. Taking advantage of iSCseq, we tried inclusive iSCseq with *in vivo* and *in vitro* scRNA-seq datasets to test whether *in vivo* OCs in the bone microenvironment are different from *in vitro* cultured OC induced by M-CSF and RANKL.

First, we built an *in vivo* single-cell bone atlas containing 74,298 cells from 32 public datasets of nine papers^29-38^ (Figure S4A and S4H). The bone cell atlas recapitulated the bone microenvironment consisting of 21 cell types, including whole hematopoietic lineage cells with hematopoietic stem cells and mesenchymal cells (Figure S4B and S4N). The bone cell atlas can provide global differentiation pathways based on RNA velocity^24,39^ (Figure S4C and S4M).

Next, we combined *in vitro* scRNA-seq datasets, including iSCseq, and the *in vivo* bone atlas (Figure 4A, S5 and S6). The area containing OCs enclosed by a two-pointed square (Figure 4A) was enlarged (Figure 4E). We shadowed the iSCseq spots inclusive *of in vivo* UMAP (Figure 4B). In the OC-focused region, *the in vitro* components were divided into two parts and had a bifurcation of the differentiation pathway (Figure 4C). One of the two parts (common OC) intersected with an *in vivo* monocyte-abundant lineage (Figure 4D, 4E and S7B). iSCseq components in iSCseq OC clusters 0 and 1 were divided as follows: *in vitro* OC points in the purple dashed line and common OC points in the orange dotted dashed line (Figure 4F). No *in vivo* bone atlas OCs were included in the *in vitro* OC zone (Figure 4G). *In vitro* OC components tended to exist on the upper right differentiation side in iSCseq alone UMAP (Figure 4H).

**Figure 4.**
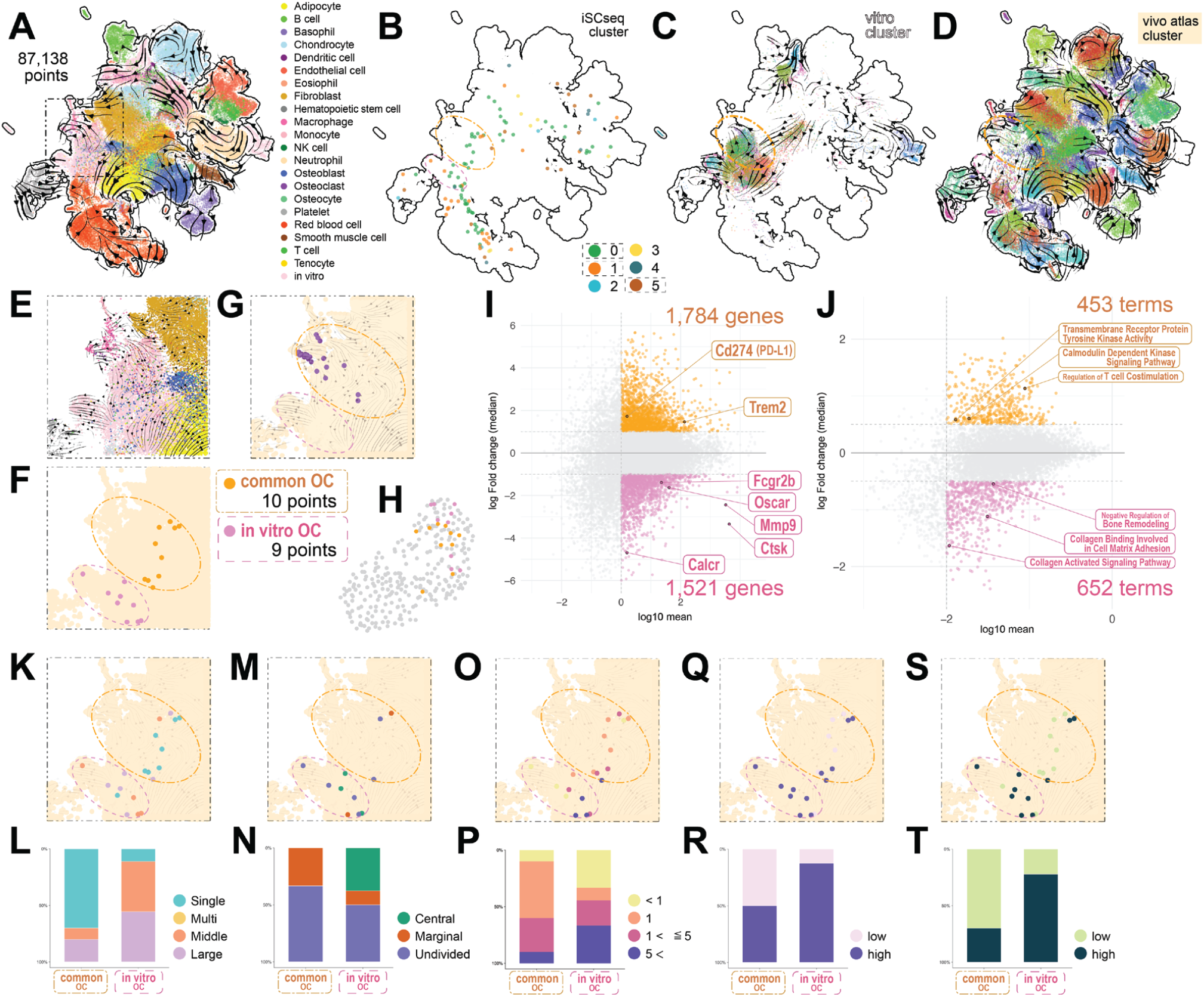
Inclusive *in vivo* iSCseq with single cell bone atlas. (A) UMAP of inclusive *in vitro* and *in vivo* iSCseq with the bone milieu atlas, which contained iSCseq spots, *in vitro* scRNA-seq cells, and *in vivo* single-cell bone atlas cells colored with cell type annotation of the *in vivo* bone atlas (Figure S4) and RNA velocity stream. *In vitro* spots, including iSCseq, are colored sakura. (B) iSCseq cluster information presentation on the inclusive *in vivo* iSCseq UMAP. The two areas where spots in iSCseq alone OC clusters accumulated are enclosed by a purple dashed circle (*in vitro* OC points) and an orange dotted dashed circle (common OC points). (C) *In vitro* iSCseq and scRNA-seq spots shown in Figure 2A with cluster numbers and scVelo streamlines. (D) *In vivo* bone atlas cells in Figure S4A with cluster numbers and scVelo streamlines. (E) Enlarged view of the area surrounded by the double-dotted line square in Panel A. (F) Two groups of OC spots in iSCseq alone clusters 0 and 1 in Figure 1E in enlarged view. (G) Spots of OC annotated by Cell Assign in the *in vivo* bone atlas in the enlarged view with scVelo streamlines and circles shown in Panel B. (H) Feature plot of the two types of OCs defined in Panel F on iSCseq alone UMAP (Figure 1E). (I) MA plot of differentially expressed genes between the two types in Panel F. (J) MA plot of differentially regulated gene ontology terms between the two OC types. (K-T) Enlarged feature plots (K, M, O, Q, S) and cumulative bar plots (L, N, P, R, T) of the basic characteristics discussed in Figure 2, 3 and S3 in the double dotted square in Panel A focused on two types of OCs. (K, L) Nuclear Rank: bin, the number of nuclei in the origin of the cell. Large, 20 or more nuclei; Middle, 5 to 19 nuclei; Multi, 2 to 4 nuclei; Single, 1. (M, N) Positional information, Central or Peripheral or Undivided in OC. (O, P) The binned number of nuclei in each sample. (Q-T) High- or low-intensity mitochondria (Q, R), and calcium (S, T) from the sampling images.

We described the characteristics between two types of OCs. Differentially expressed gene (DEG) analysis showed higher expression of *Ctsk* and *Mmp9* in *in vitro* OCs, whereas the expression of immunoreceptor tyrosine-based activation motif (ITAM) adaptor associated receptors was separated (Figure 4I). Collagen-activated pathways were activated in *in vitro* OCs, whereas ITAM-related kinase activity and T cell costimulation were upregulated in common OCs (Figure 4J). The morphological, physical, and physiological features of the two types were also evaluated (Figure 4K-T). *In vitro* OCs were mostly middle or large OCs (Figure 4K and 4L). *In vitro* OCs had only central nuclei components (Figure 4K and 4L) and a variety of components, from mononuclear to multinuclear (Figure 4O and 4P). Mitochondrial and Calcium intensities were higher *in in vitro* OCs (Figure 4Q-T).

Inclusive iSCseq revealed new subsets of OC, whose differences could not be detected using only *in vitro* assays, based on the distribution of information on large datasets and stream of RNA velocity. The iSCseq points clarified the objects to be compared and provided a larger number of DEGs compared with chromium analysis alone (Figure 1D). The new subsets were characterized by multiple pieces of evidence.

### Inclusive iSCseq with human bone tumors

*In vitro* OCs with increased acid production did not overlap with *in vivo* OC-annotated cells in physiological bones. We hypothesized that *in vitro* OC might exist under pathological conditions and be involved in excessive bone destruction. We attempted to identify subsets of OCs in the tumor microenvironment (TME), in which excessive bone resorption often causes pain and pathological fractures.

First, we performed a reanalysis of human scRNA-seq public datasets from giant cell tumor (GCT) of bone in four patients^40^ (Figure S8), osteosarcoma in six patients^41^ (Figure S9), bone metastasis of breast cancer in two patients^42^ and prostate cancer in nine patients^43^ (Figure S10). Malignant cells were identified by copy number variant (CNV) (Figure S8F, S8G, S9F, S9G, S10F and S10G). Osteoclasts in the GCT of bone and prostate cancer metastasis were all normal; instead, some OCs in osteosarcoma and all OCs in breast cancer were malignant (Figure S8N, S9N and S10N).

We then performed the final inclusive iSCseq with all datasets shown in the previous main figures (Figure 5A, 5D and S11). *In vivo* wild-type cell types in the bone atlas were well separated on the UMAP and could be indicators for cell type identification (Figure 5B). Blood cells on the upper side were almost normal based on the CNV (Figure 5C).

**Figure 5.**
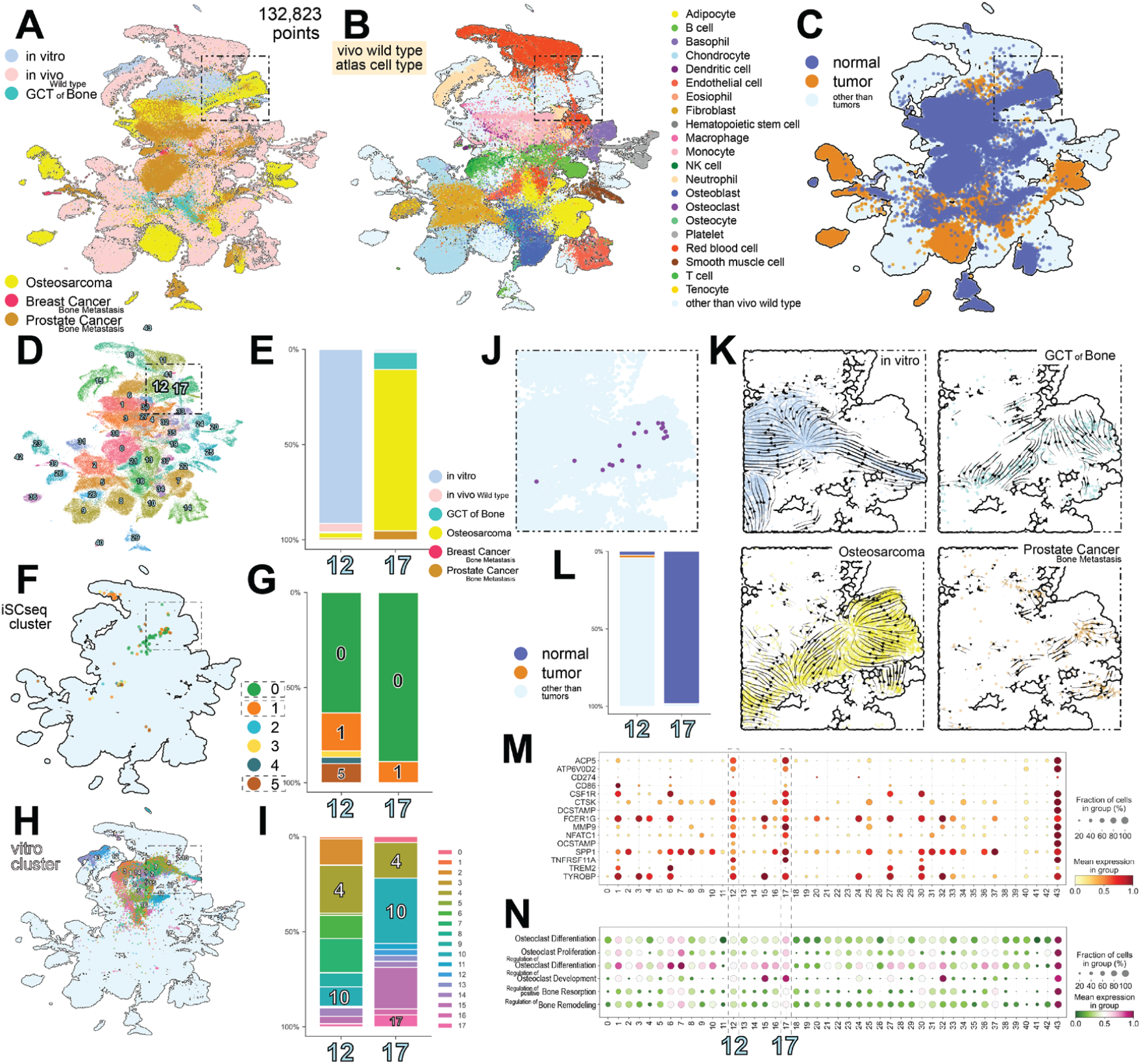
Final inclusive iSCseq with human bone tumors and metastases. (A) UMAP of the final inclusive iSCseq with the bone milieu atlas, which contained iSCseq spots and humanized inclusive *in vitro* iSCseq cells (*in vitro*), humanized *in vivo* bone atlas cells (*in vivo* wild type), and four types of human tumors and bone metastases. *In vitro* OC-enriched region is enclosed by a two-dot chain square. (B) Cell type annotation in the *in vivo* wild type bone atlas. (C) Normal or tumor information based on CNV score for each tumor or bone metastasis. (D) Cluster information presentation on the UMAP. OC-enriched region is enclosed by a two-dot chain square. (E) Cumulative bar plot of samples in OC-enriched clusters 12 and 17 in final inclusive iSCseq. (F, G) iSCseq alone spots with cluster information in Figure 1E on UMAP (F) and cumulative bar plot in clusters 12 and 17 (G). (H, I) *In vitro* spots with cluster information in Figure 2A on UMAP (H) and cumulative bar plot in clusters 12 and 17 (I). (L) *In vivo* bone atlas OCs (purple) in the enlarged view of a two-dot chain square in Panel A. (M) Enlarged view of a two-dot chain square by sample category with scVelo streamlines. (N) Cumulative bar plot of normal or tumor cells evaluated by copy number variant in clusters 12 and 17. (O) Dot plot of OC-related and immune-tyrosine associated motifs-related gene expression. (P) Dot plot of gene ontology terms related to OC.

We shadowed the iSCseq spots (Figure 5F) and inclusive *in vitro* iSCseq spots (Figure 5H) on UMAP. Components of *in vitro* OC-enriched clusters (4, 10 and 17) were enriched in a double-dotted square (Figure 5H). *In vitro* OC was distributed on the right side of the square (Figure S11E). This square also contained *in vivo* bone OCs (Figure 5J).

The main clusters in the square were clusters 12 and 17 in the final inclusive iSCseq analysis (Figure 5D). Cluster 12 contained abundant *in vitro* components. Instead, cluster 17 consisted of normal cells from GCT of bone, osteosarcoma and prostate cancer bone metastasis added to iSCseq spots, including *in vitro* OCs defined in Figure 4F (Figure 5E, 5L,5K and S11E). Cluster 17 contained consistently more differentiated OCs in iSCseq alone (Figure 5G) and inclusive *in vitro* iSCseq (Figure 5I) than cluster 12. Clusters 12 and 17 exhibited an osteoclast phenotype in terms of gene expression (Figure 5M) and ontology (Figure 5N). Morphological and physiological properties of iSCseq spots were summarized in Figure S12B-F. From the RNA velocity analysis of each sample category (Figure 5K and S12A), *in vitro* spots and tumor cells overlapped, but had opposite relative directions of differentiation.

We found that *in vitro* OC intersected with cluster 17, which is abundant in OC in tumors and bone metastasis. However, OCs in the tumor microenvironment were derived from another origin different from *that of in vitro* monocytes based on RNA velocity analysis. Different signaling intensities of CSF1 and RANKL and the transcriptional factor activity of NFATc1 and IRF8 may affect the distribution in cluster 17 (Figure S11G, S11I and S11M).

### DNA methylome for sampled nuclei

Epigenetic control of transcription is important in the OC differentiation process^44,45^. DNA methylation is an epigenetic regulator that has been extensively investigated in developmental and cancer biology.

We performed whole genome bisulfite sequencing (WGBS) to examine the DNA methylation levels of one or a few nuclei using seven samples with the same sampling method as iSCseq (Figure S12A, S12B and S12C). WGBS showed different methylation patterns even in the same OC (Figure S12D). A marked difference in DNA methylation levels was observed, particularly in the promoter region (Figure S12E and S12F). This is the first evidence of DNA methylation levels in the same multinucleated cells.

### Centrality-focused network analysis of cellular components

iSCseq revealed that the terminal cluster in the OC differentiation pathway consisted not only of multinuclear cells but also of small ordinary cells. Next, the fusion process is discussed separately from the differentiation process. iSCseq identifies the origin of cells in each sample, thus enabling the examination of the connections between cellular components.

First, we counted the frequency of occurrence of iSCseq cluster numbers based on cellular identity from live cell images and summarized them into a colocalization map on the corresponding UMAP. Then, page rank and closeness centrality extracted the core clusters of connection by the OC fusion step according to the Nuclear Rank defined in Figure 2I and 2J from Single to Large (Figure 6A and 6B). Next, we separately discussed the connection to terminal cluster 10 of inclusive *in vitro* iSCseq (Figure 6C). Interestingly, cluster 10 was connected to cluster 13 in the same single cell. Cluster 13 was classified as neutrophil by automatic cell annotation *in vitro* (Figure 2E) and *in vivo* (Figure 4A and S7A). In the multi nuclei phase (two to four), components apart from the differentiation pathway were involved in fusion. In the middle and large phases (five or more nuclei), the connection within cluster 10 was noticeable. Clusters 4 and 17 in the OC differentiation pathway were also involved in fusion, but variations in connection were fewer than those in clusters 10 and 13 (Figure S14A-C).

**Figure 6.**
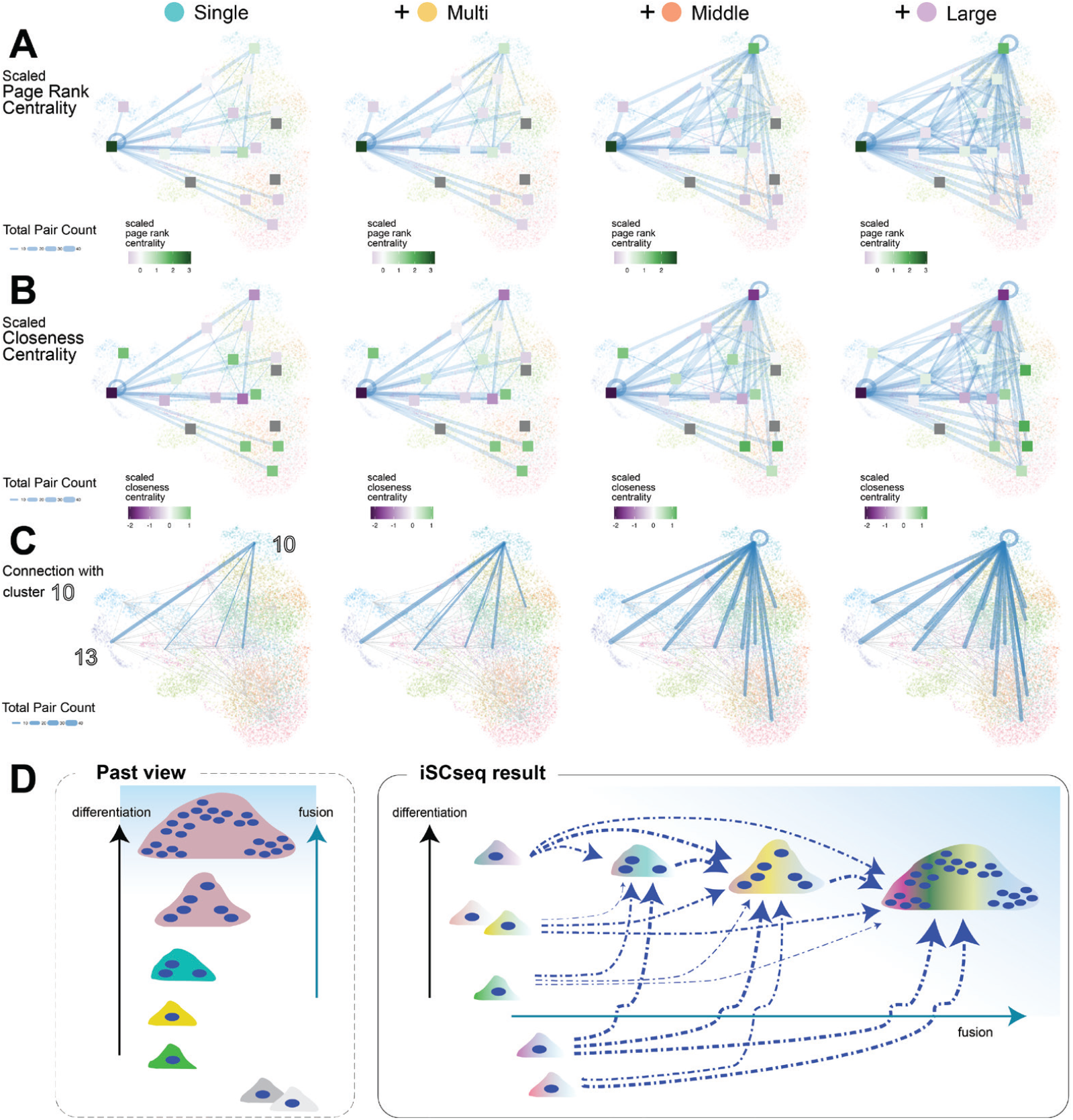
Colocalization of iSCseq components with centrality analysis. (A, B) Page rank (A) and closeness (B) centrality analysis with linkage of iSCseq components on inclusive *in vitro* iSCseq UMAP (Figure 2A) by gradually increasing Nuclear Rank. Colocalization of iSCseq components between different clusters as a node and within the same cluster as a self-loop. The edge or loop linewidth reflects the total number of pairs (10 or more highlighted). Nuclear Rank: bin, the number of nuclei in the origin of the cell. Large, 20 or more nuclei; Middle, 5 to 19 nuclei; Multi, 2 to 4 nuclei; Single, 1. (C) Edges and self-loops connected to the inclusive *in vitro* iSCseq terminal cluster 10. (D) Schematic representation of the osteoclast differentiation and fusion process. The single-pointed arrows represent the fusion order, and the linewidth reflects the contribution of each fusion stage.

iSCseq allowed us to discuss the components involved in fusion by differentiation stage (Figure 6D). Differentiation and fusion have been discussed without any cell-by-cell evidence of multinucleated cells. iSCseq showed that neutrophils unrelated to the OC differentiation pathway were involved in early cell fusion. In the later fusion phase, not only cells in the differentiation pathway but also more variegated cells were recruited. The same centrality-focused network analysis for iSCseq alone showed that terminal cluster 0 was the core node of connection with all other clusters (Figure S15).

## Discussion

Traditional single-cell technology using microfluid devices has clarified the cellular heterogeneity in terms of gene expression^46^. However, there is a lack of cellular identity as a trade-off for achieving a high throughput. Moreover, scRNA-seq separates the differences with the cell as the smallest unit. The subcellular components were beyond the scope of scRNA-seq. Our iSCseq approach is the first realistic proof-of-concept approach for detecting subcellular heterogeneity within a single cell considering morphological and physiological annotations.

Inclusive analyses of iSCseq connected conventional scRNA-seq to the cellular or subcellular reality. The origin of the cells or subcellular positions inside a cell can be determined using this combinatorial approach. Large number of scRNA-seq datasets compensate throuputness in iSCseq approach. In contrast, we observed a higher number of genes using iSCseq spots compared to traditional scRNA-seq (Figure 1D). For example, cross-species analysis limits the number of genes in the process of homologue conversion. However, comparison between the focused iSCseq points extracted from high-resolution UMAP allowed the comparison of not only a larger number of genes, but also morphological and physiological features with multi-layered evidence. Cost reduction of sampling and plate-based sequencing is key to random unbiased sampling and sequencing to solve biological and clinical questions. Because scRNA-seq and iSCseq have complementary effects, we recommend using iSCseq in combination to increase the resolution and gain biological meaning of subcellular components. iSCseq can be extended to other omics approaches such as DNA methylome (Figure S13). Single-cell technology can be augmented using iSCseq.

Multinucleated or large cells are beyond the scope of the trendy single-cell approach. However, iSCseq enabled us to discuss heterogeneity within giant cell populations, which has been difficult to analyze. iSCseq clarified that not all cellular components in *in vitro* OC culture were involved in the OC differentiation pathway, and some undifferentiated components were caught or engulfed in the OC fusion process. Trendy trajectory analysis tends to instill the preconception that differentiation proceeds in a single direction. The iSCseq approach provides insights into a complex differentiation paradigm involving cell fusion.

Differentiation and fusion have been confusingly discussed without any cell-by-cell evidence of multinucleation. iSCseq revealed cellular identity with multilayered evidence and allowed us to discuss differentiation and fusion separately for the first time. Centrality is a concept commonly used in modern information science, such as in website network analysis^47^. We examined two different types of centralities, page rank^48^ and closeness^49^, to discuss the characteristic fusion process in multinucleated cells and found that cells such as neutrophils, which are not related to the OC differentiation pathway, were involved in the early OC fusion process. Our differentiation and fusion model provided the first insights into the complex fusion process acquiring subcellular diversity with quantitative evidence. This model may explain why bone marrow cells, which contain diverse cell populations rather than sorted monocytes, have been frequently used in *in vitro* osteoclast differentiation experiments. iSCseq is a promising toolkit to investigate subcellular community to enhance cell physiology.

Existence of OC subsets is a hot topic in the field of skeletal biology. Recently, pathological OC from a different cell source, AtoM, has recently been proposed^50^. Our findings depicted two subsets *in vitro*, and one subset was common in physiological bones. *In vitro* OC promoted higher OC proliferation and regulation of bone resorption and existed in the tumor microenvironment (TME). OCs in the TME have another source from *in vitro* assays based on inclusive iSCseq with RNA velocity. Our inclusive analyses, which connected large datasets with iSCseq, captured new subsets depending on the environment with multilayered evidence of morphology and physiology. The definition of OC can be further clarified in the omics era.

Furthermore, inclusive iSCseq identified candidate switching molecules between subsets of OCs. Epigenetic controls by the transcriptional factors NFATc1 and IRF8 are important in OC differentiation^44,45^. At single-cell resolution, IRF8 activity was identified as a candidate regulatory transcriptional factor between *in vitro* OC and common OC (Figure S5I).

In addition, analyses of differentially expressed genes showed that immunoreceptor tyrosine-based activation motifs (ITAMs) and related costimulatory factors may function as separators between the two OC subsets (Figure 4I). ITAMs and costimulatory factors were believed as the third OC differentiation factor from the experiments using knockout or knockdown mice^12,51,52^. Based on the results of differentially expressed genes and terms (Figure 4I, 4J and 5K), plasmalemmal molecules, such as receptors of costimulatory signals, are therapeutic candidate targets for bone resorption. Immunomodulatory drugs targeting Fc receptors^52,53^ and TREM2^54,55^ coupled with TYROBP/DAP12 may be good targets for osteoporosis treatment in physiological bones. iSCseq demonstrated for the first time the physiological and pathological roles of ITAMs and related costimulatory signals.

In conclusion, a newly developed method, iSCseq, demonstrated subcellular heterogeneity of gene expression and its biological significance. iSCseq proved that multiple differentiation stages can be embedded in a cell. The physical responses and physiological distribution of mitochondria and calcium were confirmed using gene expression analysis. Inclusive iSCseq with similar datasets elucidated the physiological mechanisms and human skeletal pathogenesis in osteoimmune diversity with multi-layered evidence. Centrality-focused network analysis expounded the diversity acquisition process in multinucleation and the hidden key players in the complex fusion process. iSCseq provided clues that could not be obtained using conventional research methods. iSCseq is expected to be the next core technology in cell biology.

## STAR ⋆ METHODS

Detailed methods are provided as the following:

- KEY RESOURCE TABLE
- RESOURCE AVAILABILITY
  - Lead contact
  - Data availability
  - Supplemental resources
  - Ethical considerations
- EXPERIMENTAL MODELS
  - Mice
  - Primary bone marrow cell cultures and in vitro osteoclast formation assay
- METHODS DETAILS
  - Live cell staining
  - Sampling of intracellular components
  - Library construction for intra-single cell RNA-seq with micro input
  - Library construction from sampled nuclei for whole genome bisulfite sequencing (WGBS)
  - Next generation sequencing of RNA-seq libraries
- QUANTIFICATION AND STATISTICAL ANALYSIS
  - Bioinformatical analysis of intra-single cell RNA-seq (iSCseq)
  - Public resource of scRNA-seq datasets
  - Analysis of images
  - Bioinformatical analysis of DNA methylation by whole genome bisulfite sequencing
  - Network and centrality analysis of iSCseq components
  - Statistics

## ACKNOWLEDGEMENT

Live cell imaging and intracellular sampling were technically supported by Yokogawa Electric Corporation during the planning stage of the experiments. Our next generation sequencing was successful with a highly elaborate manufacturer’s kit from Takara Bio and Clontech. The supercomputer SHIROKANE at the Human Genome Center at the University of Tokyo was partially used in the calculation. The platform for Advanced Genome Science in Japan supported this project of elaborate RNA-seq. NGS Expo Osaka 2022 offered part of whole genome sequencing as the best presentation award. NGS Expo Osaka 2022 offered part of whole genome sequencing as the best presentation award. Kashiwa sequencing team and NGS core facility at the Research Institute for Microbial Diseases of Osaka University supported sequencing.

This work was funded by JSPS KAKENHI grant numbers JP 20K22973, 21K16703, 22KK0272, 23K15736 (HO), JP 21K19589 (HH and HO), JP 21K19883 (UC, HH and HO), JP 22K10212 (SK, HH and HO), JP 16H06279 (HO, PAGS: Platform for Advanced Genome Science) and 22H04925 (HO, PAGS ver.2), AMED grant number JP21zf0127002 (HH and HO) and JP21bm0704071 (HH, MS and HO), “Creation of fully circular bio-adaptive materials driven by bio-polymer big data” Material DX (HH and HO), KAWAI Foundation for Sound Technology and Music 2021 (HO), Takeda Science Foundation 2022 (HO), Mochida Memorial Foundation for Medical and Pharmaceutical Research 2022 (HO), Astellas Foundation for Research on Metabolic Disorder 2023 (HO) and FY2020 Invitational Fellowships for Research in Japan L20539 (RB).

## AUTOHR CONTRIBUTIONS

Conceptualization: HO, MH, SK, MH, HK, KO, RB, S. Tanaka

Methodology: HO, YT, YO, MS, HK, YS, HH

Investigation: HO, YT, YO, S. Tani, DM, JM, KM, SO, FY

Visualization: HO, YT, S. Tani, AT, JM

Funding acquisition: HO, SK, DM, TS, YS, S. Tanaka, UC, HH

Project administration: HO, HH

Supervision: RB, FG, TS, YS, S. Tanaka, HH

Writing – original draft: HO

Writing – review & editing: HO, AT, S. Tanaka, RB, UC, HH

## INCLUSION AND DIVERSITY

We supported inclusive, diverse, and equitable conduct of research.

## Supplemental figures

**Figure S1.**
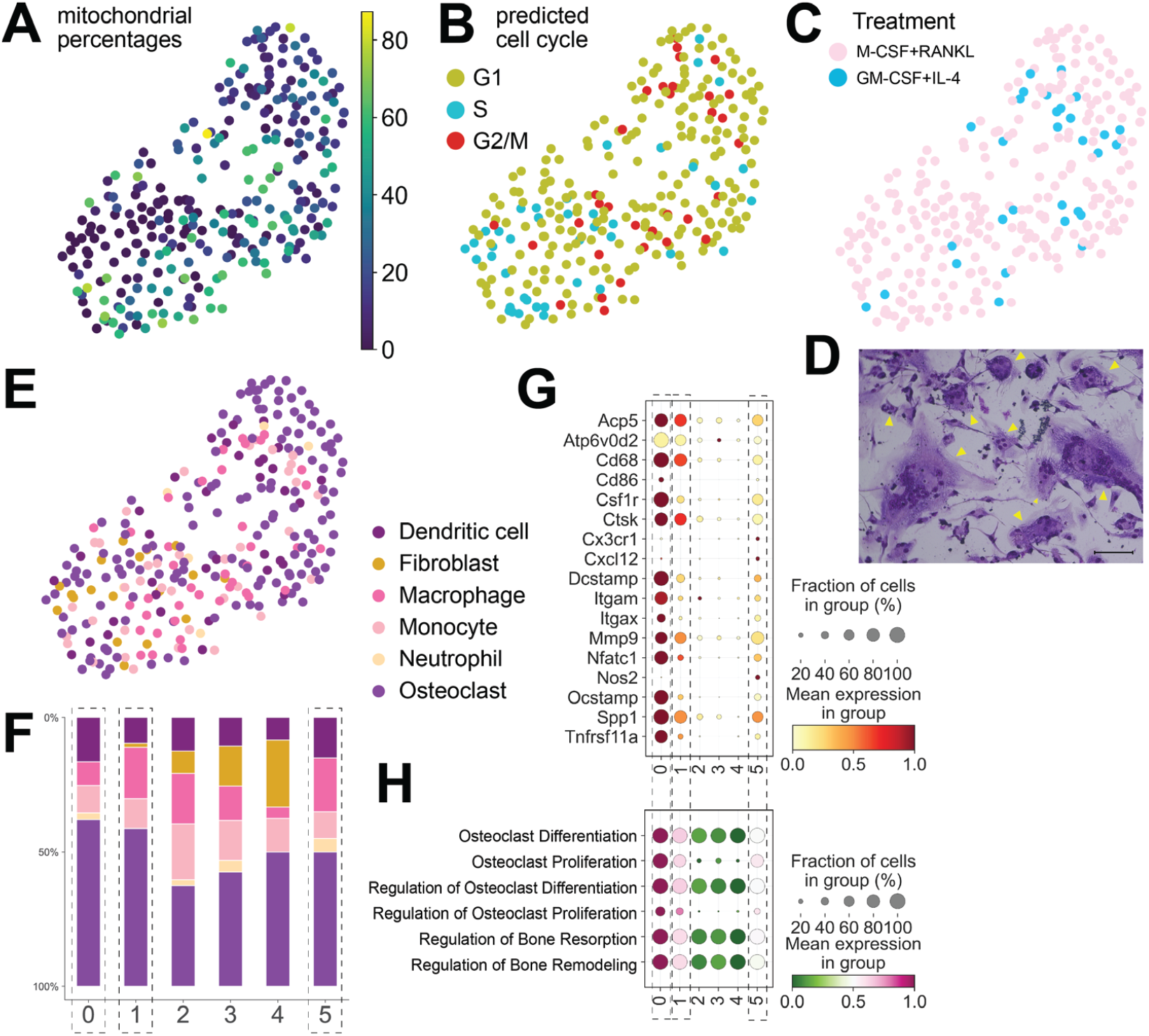
Basic parameters of intra-single cell sequencing (iSCseq) alone. (A-C) Feature Plots of mitochondrial percentage (A) and of cell cycles (B) of inductive treatment options (C) from iSCseq. (D) Representative foreign body giant cells (FBGCs) stained with May-Grünwald Giemsa. Yellow arrowheads indicate FBGC. Bar = 50 μm. (E, F) Automatic cell-type annotation using CellAssign on feature plot (E) and cumulative bar plot (F). Osteoclast (OC)-enriched clusters are enclosed in dashed rectangles. (G, H) Dot plots of OC-related genes (G) and GO terms (H). OC-enriched clusters are highlighted with dotted rectangles.

**Figure S2.**
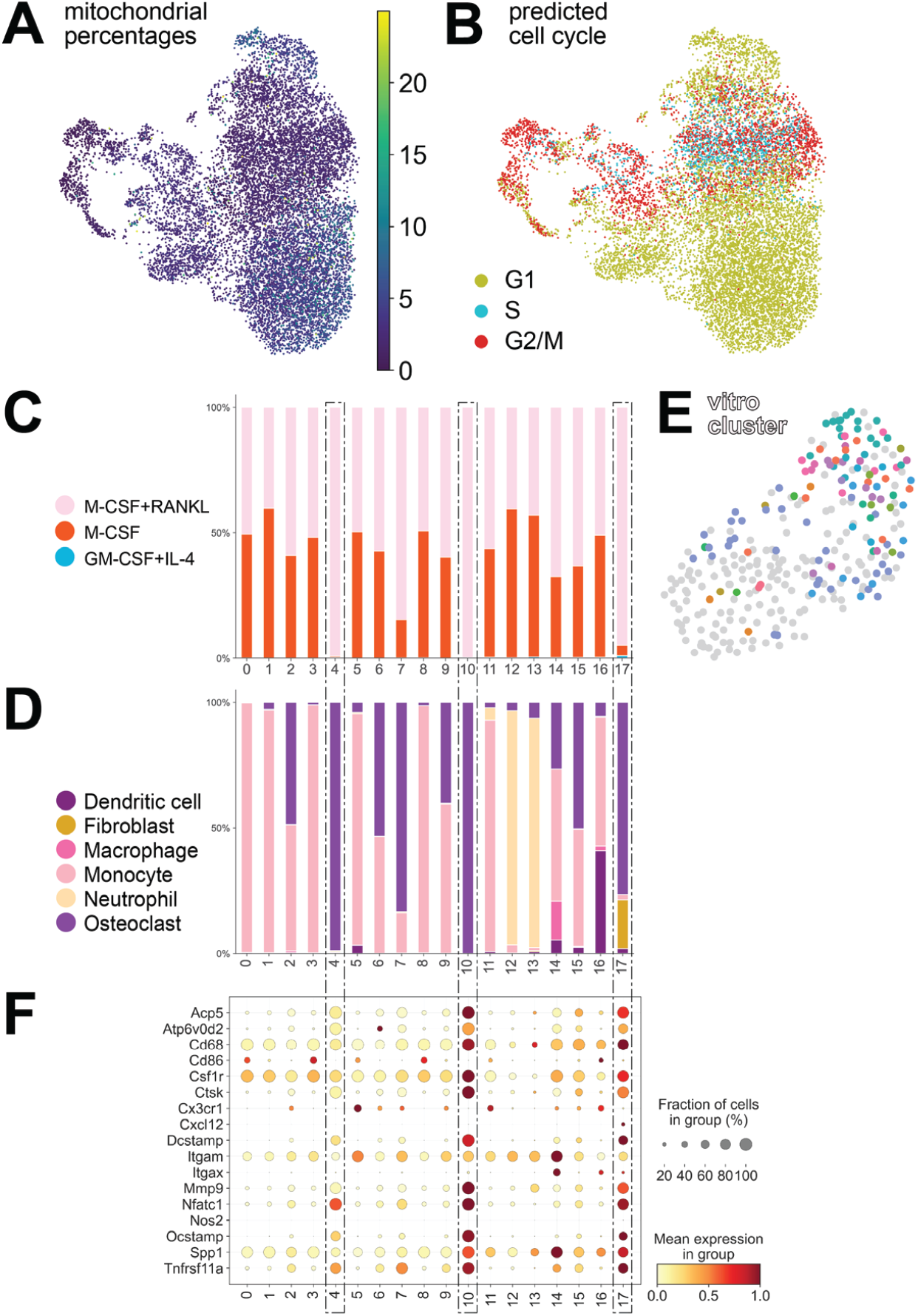
Basic parameters of inclusive iSCseq with *in vitro* cultured scRNA-seq datasets. (A, B) Feature Plots of mitochondrial percentage (A) and of cell cycle (B) from inclusive *in vitro* iSCseq. (C, D) Cumulative bar plot of cytokine treatment (C) and of the cell type using CellAssign (D). (E) Inclusive *in vitro* iSCseq cluster information spotted on iSCseq alone UMAP. (F) Dot plot of osteoclast-related genes. Osteoclast-enriched clusters are enclosed by dotted dashed rectangles.

**Figure S3.**
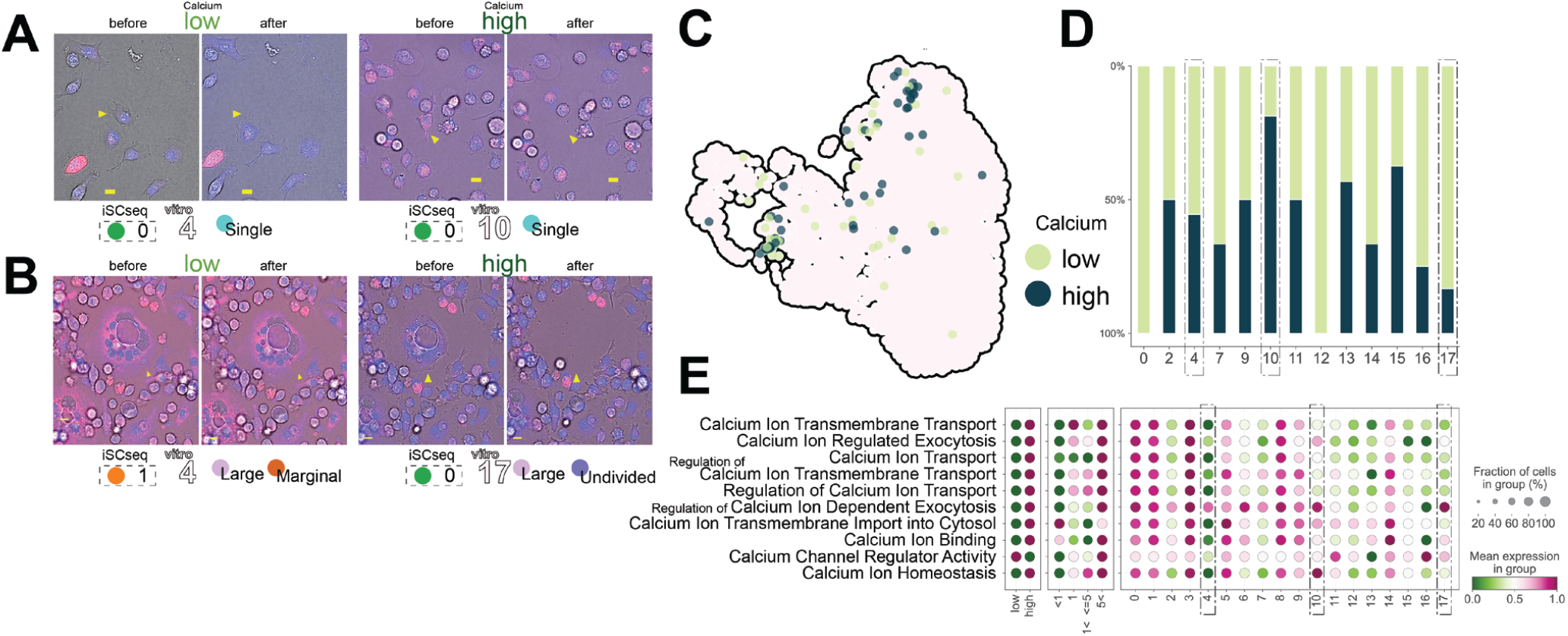
Morphophysiological analysis of calcium with the transcriptome. (A, B) Representative sampling cases of single nucleated cells (A) and multinucleated OC (B) with qualitatively high or low dichotomized intensity of calcium (Ca) with inclusive *in vitro* iSCseq cluster information in Figure 2A. Blue: Nuclei stained with Hoechst 33342; Red: CalbryteTM 590 ABD. Bar = 10 μm. (C, D) High or low dichotomized intensity of Ca staining in each iSCseq sample. (C) Feature plot of iSCseq spots with high or low Ca intensity on inclusive *in vitro* iSCseq UMAP. (D) Cumulative bar pot of high or low Ca intensity in each cluster. (E) Dot plot of the top 10 gene ontology (GO) terms including Ca using decoupleR and AUCell by high or low Ca intensity from images (left), by inclusive *in vitro* iSCseq cluster (center), and binned number of nuclei in each sample (right). OC-enriched clusters are surrounded by dotted dashed rectangles.

**Figure S4.**
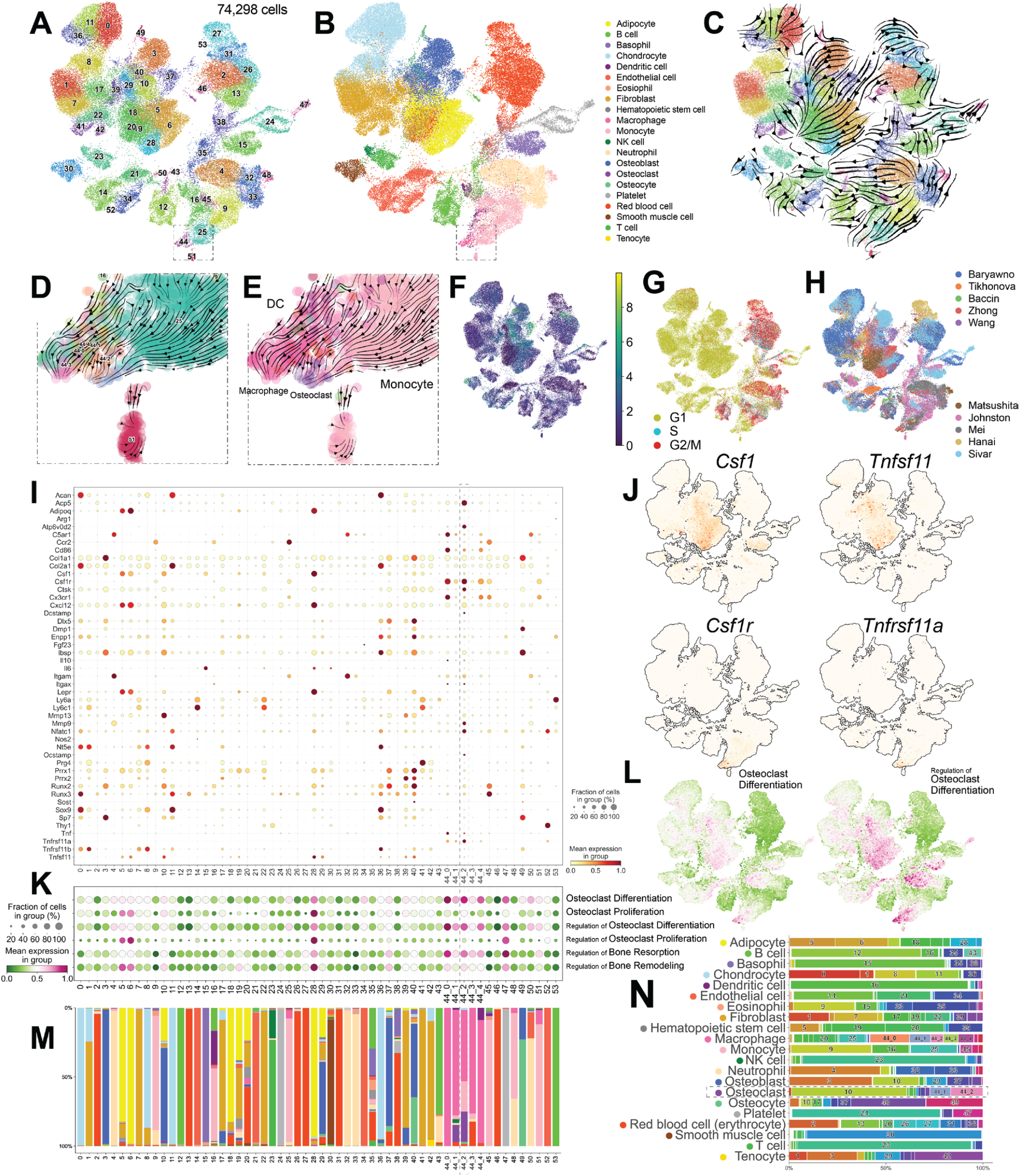
*In vivo* single cell bone atlas. (A) UMAP of the *in vivo* single-cell bone and bone marrow atlas from 32 datasets from ten papers, including wild-type or wild-type equivalent control cells. (B) Automatic cell type annotation using CellAssign. (C) scVelo RNA velocity streamlines. (D, E) Enlarged view of the area surrounded by a dotted dashed square in Panel A and B. Cluster information after subclustering osteoclast (OC)-enriched cluster 44 (D) and CellAssign cell type (E) with RNA velocity streamlines. (F-H) Feature Plots of mitochondrial percentage (F), of cell cycle (G) and of paper authors (H). (I) Dot plot of gene expressions of skeletal gene set. (J) Feature plots of gene expression related to OC differentiation factors. (K, L) Dot plot (K) and feature plot (L) of Gene ontology terms related to OC. (M) Cumulative bar plot of cell types by CellAssign by cluster. OC-enriched cluster 44_2 is enclosed by a dotted rectangle. (N) Cumulative bar plot of cell types using CellAssign. Cluster numbers including >5% of the cells in each cell type are shown.

**Figure S5.**
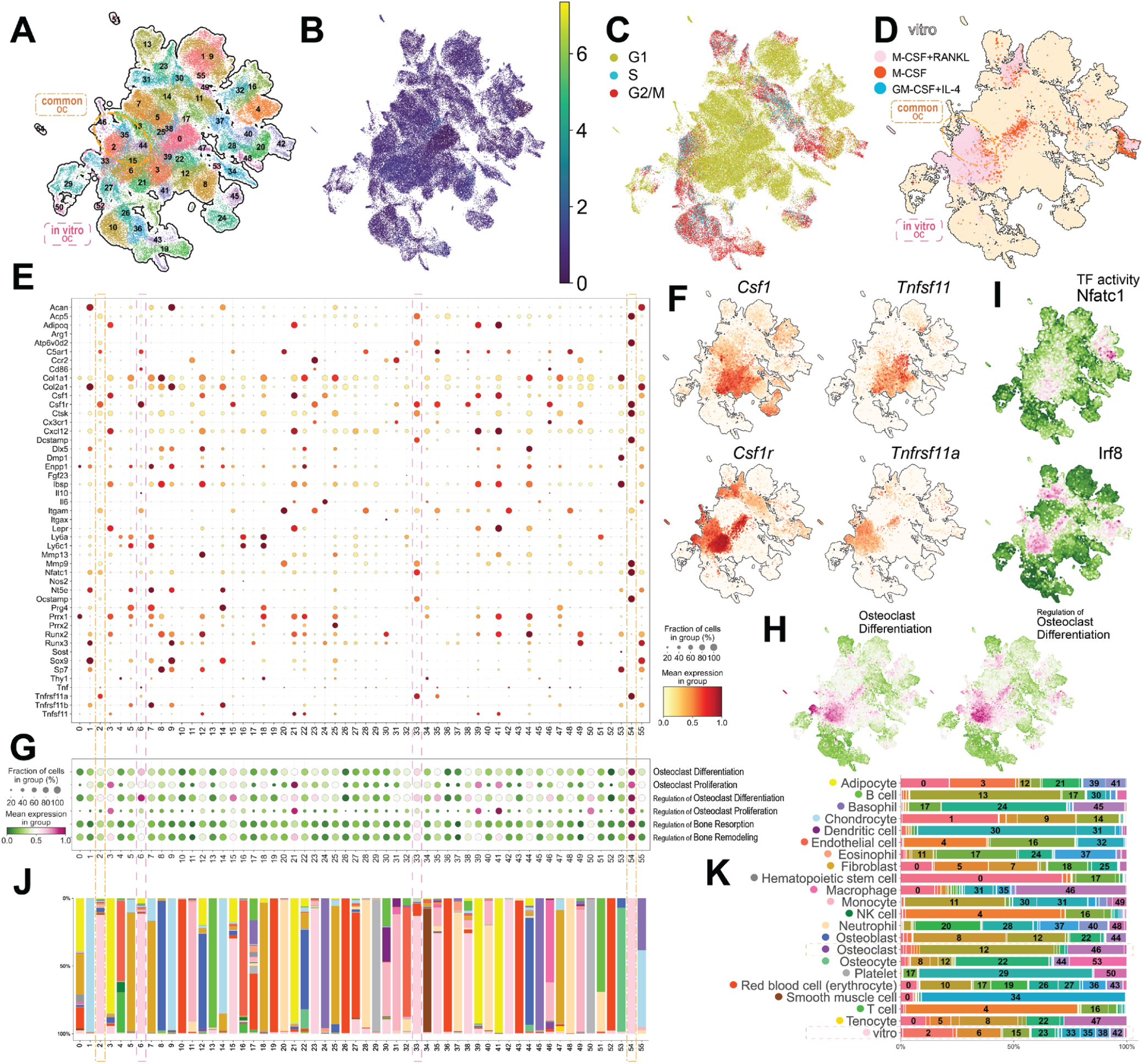
Basic parameters of inclusive *in vivo* iSCseq with single cell bone atlas (Figure 4). (A) UMAP of inclusive *in vitro* and *in vivo* iSCseq. The two groups of differentiated osteoclasts (OCs) identified by iSCseq (Figure 4F) are enclosed by a purple dashed circle (*in vitro* OC points) and an orange dotted dashed circle (common OC points). (B-D) Feature Plots of mitochondrial percentage (B), of cell cycle (C) and of treatment with cytokines for *in vitro* spots (D). (E) Dot plot of gene expression in the skeletal gene set. (F) Feature plots of gene expression related to OC differentiation factors. (G, H) Gene ontology analysis of osteoclast differentiation was performed using decoupleR and AUCell. Dot plot (G) and feature plots (H). (I) Feature plot of regulatory transcriptional factors calculated using decoupleR and AUCell. (J) Cumulative bar plot of cell types using CellAssign by cluster. Clusters including the two types of differentiated OCs (mainly defined by circles’ position on UMAP and Tnfsf11a upregulation) are enclosed by the corresponding color of rectangles in Panel A. (K) Cumulative bar plot of inclusive *in vitro* and *in vivo* iSCseq clusters. Clusters containing > 5% cells in each cell type were numbered.

**Figure S6.**
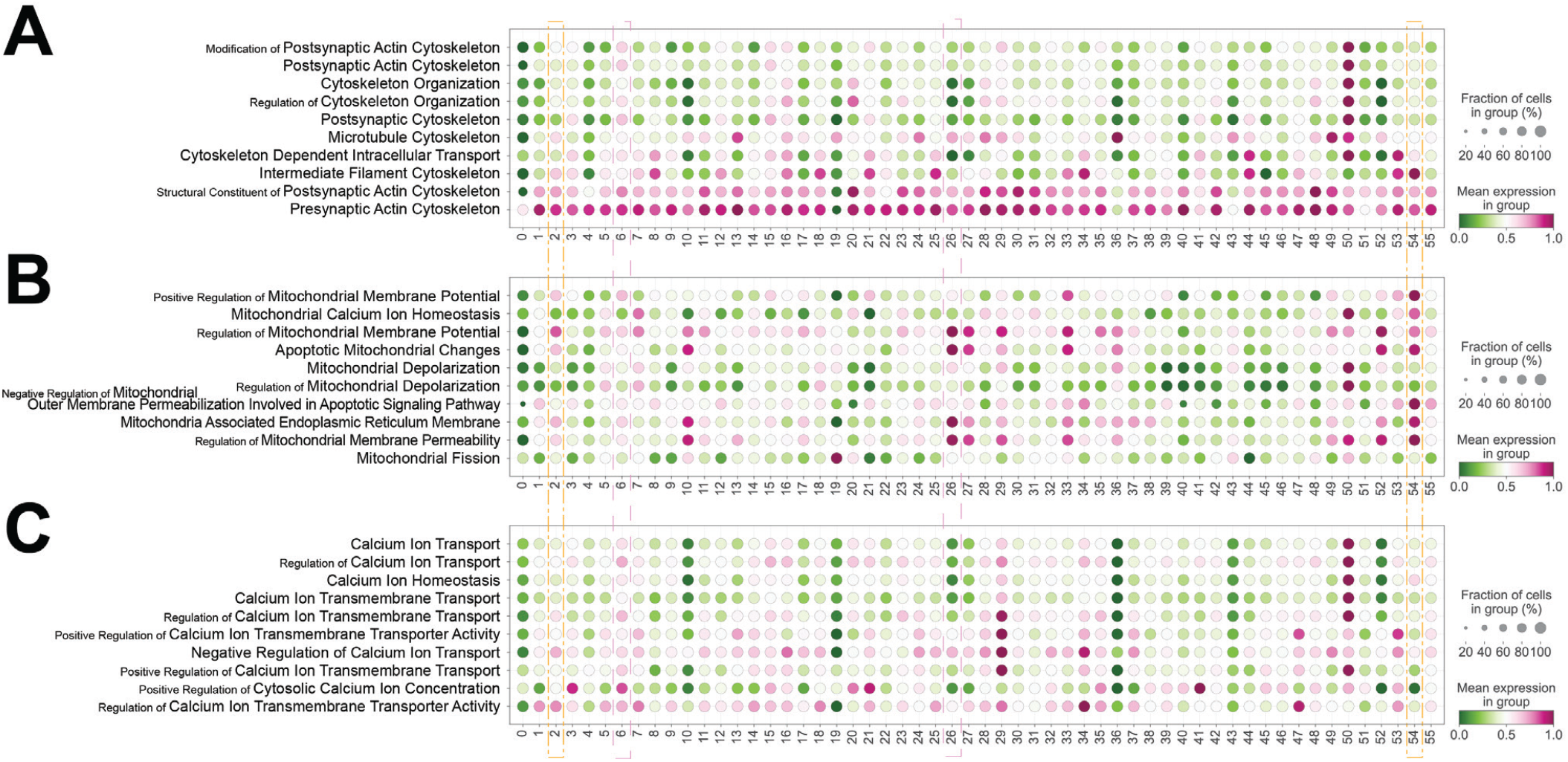
Gene ontology analysis in inclusive *in vivo* iSCseq with single cell bone atlas (Figure 4 and S5). (A-C) Dot plots of the top 10 gene ontology terms including cytoskeleton (A), mitochondria (B) and calcium (C), in inclusive *in vitro* and *in vivo* iSCseq. Clusters including the two types of differentiated OCs (defined in Figure 4F) are enclosed by the corresponding squares (*in vitro* OC, purple dashed rectangles; common OC, orange dotted dashed rectangles).

**Figure S7.**
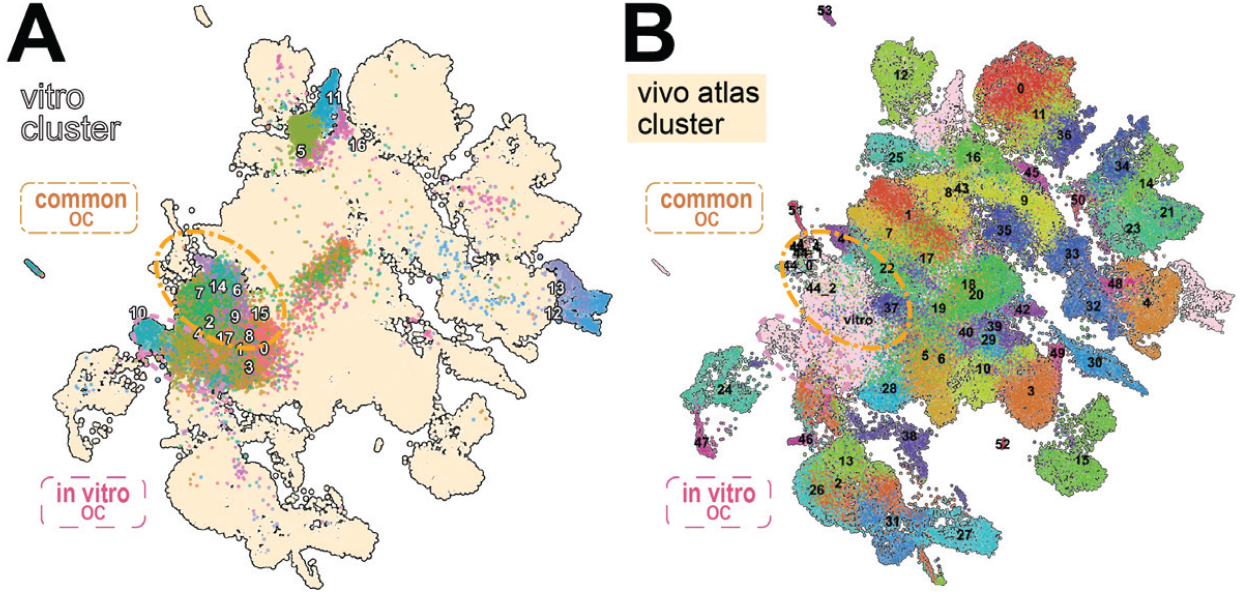
Feature plots of inclusive *in vivo* iSCseq with single cell bone atlas (Figure 4). (A) Inclusive *in vitro* iSCseq spots shown in Figure 2A with *in vitro* cluster numbers. (B) *In vivo* bone atlas cells (Figure S4A) with *in vivo* cluster numbers. a purple dashed circle represents *in vitro* OC and an orange dotted dashed circle represents common OC in Figure 4F.

**Figure S8.**
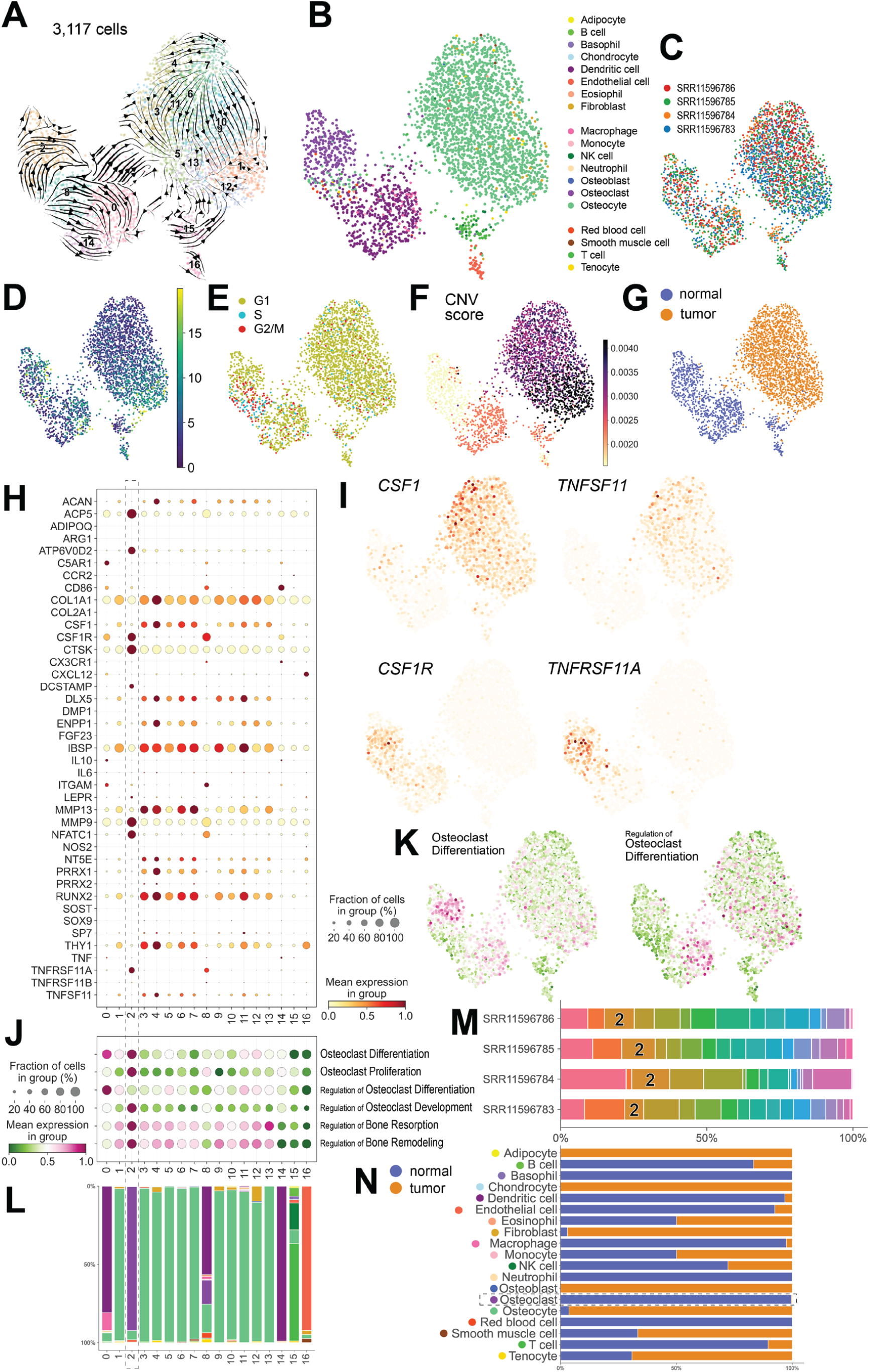
Basic parameters of giant cell tumor of bone datasets from four patients. Reanalysis of public scRNA-seq datasets from four patients with giant cell tumor (GCT) of the bone. (A) UMAP with scVelo streamlines. (B) Automatic cell type annotation using CellAssign. (C-F) Feature Plots of patient identity (C), of mitochondrial percentage (D), of cell cycle (E), of copy number variant (CNV) score calculated using infercnvpy (F) and of normal or tumor information based on CNV score (G). (H) Dot plot of gene expression in the skeletal gene set. (I) Feature plots of gene expression related to osteoclast (OC) differentiation factors. (J, K) Dot plot (J) and Feature plots (K) of OC-related terms. (L) Cumulative bar plot of cell types using CellAssign. OC-enriched cluster 2 is highlighted. (M) Cumulative bar plot of cluster components by patient. OC-enriched cluster 2 is highlighted. (N) Cumulative bar plot of normal or tumor by cell type.

**Figure S9.**
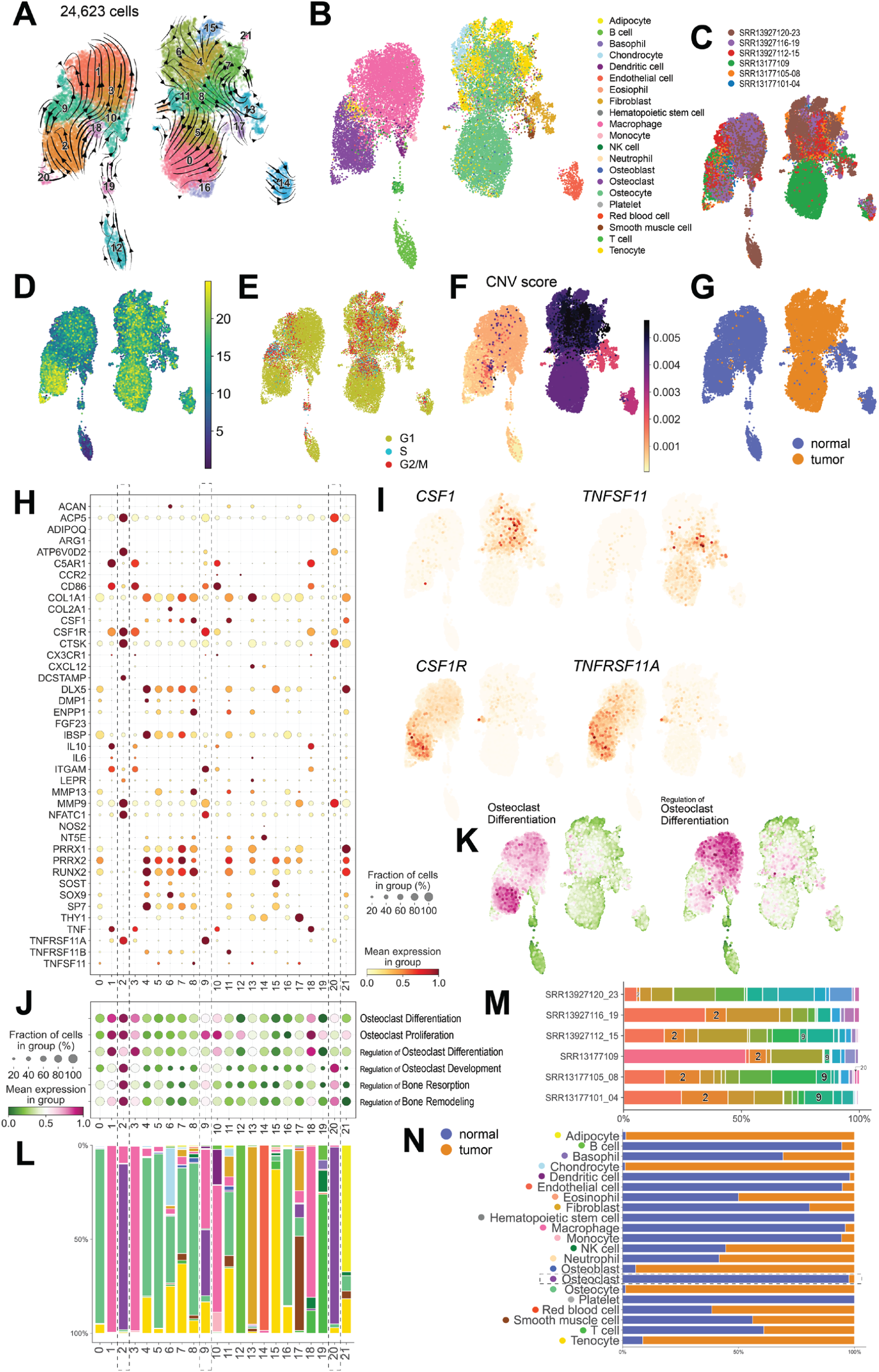
Basic parameters of osteosarcoma datasets from six patients. Reanalysis of public scRNA-seq datasets from six patients with osteosarcoma. (A) UMAP with scVelo streamlines. (B) Automatic cell type annotation using CellAssign. (C-F) Feature Plots of patient identity (C), of mitochondrial percentage (D), of cell cycle (E), of copy number variant (CNV) score calculated using infercnvpy (F) and of normal or tumor information based on CNV score (G). (H) Dot plot of gene expression in the skeletal gene set. (I) Feature plots of gene expression related to osteoclast (OC) differentiation factors. (J, K) Dot plot (J) and Feature plots (K) of OC-related terms. (L) Cumulative bar plot of cell types using CellAssign. OC-enriched clusters 2, 9 and 20 are highlighted. (M) Cumulative bar plot of cluster components by patient. (N) Cumulative bar plot of normal or tumor by cell type.

**Figure S10.**
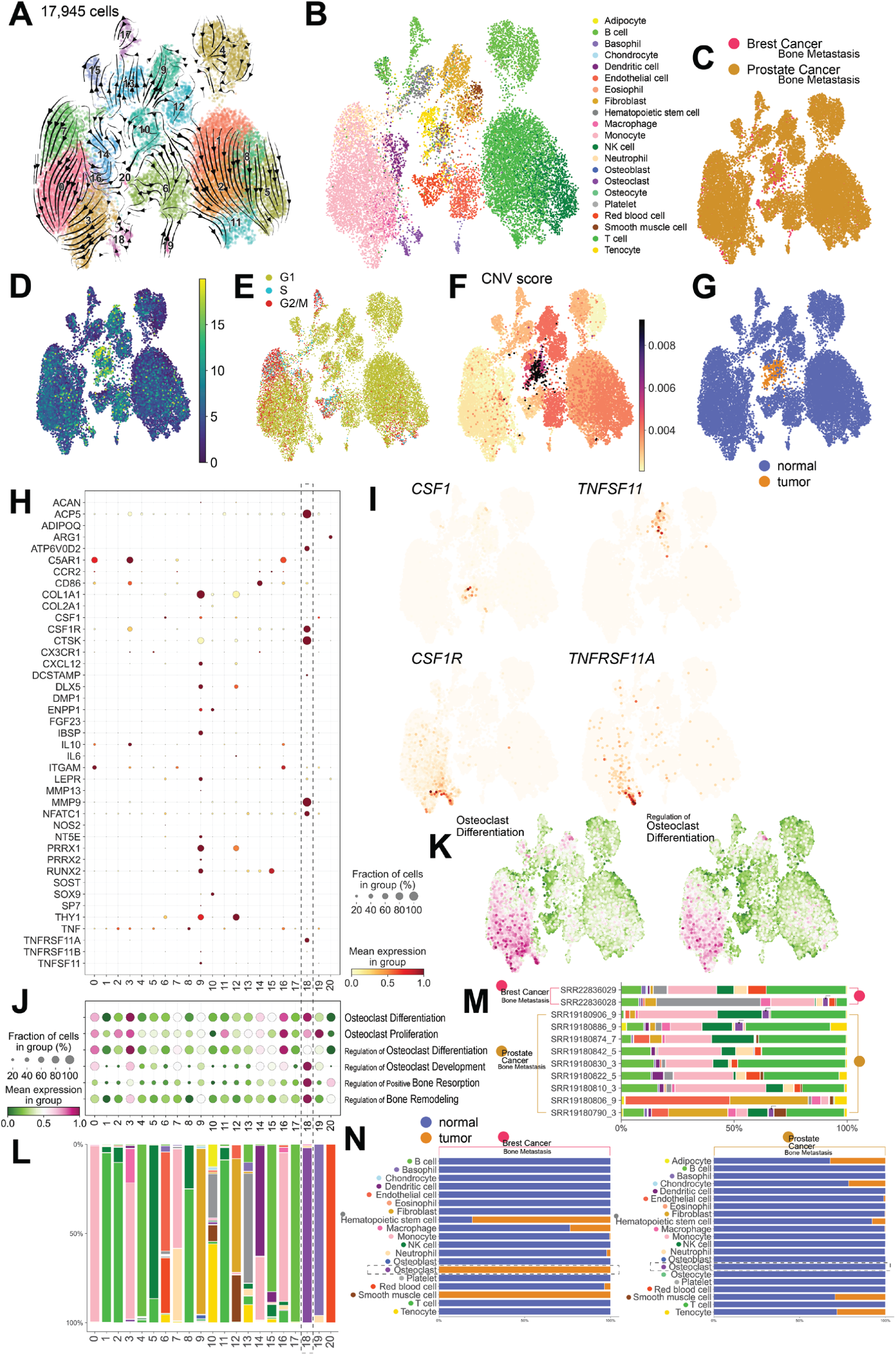
Basic parameters of bone metastasis of breast and prostate cancer. Reanalysis of public scRNA-seq datasets of bone metastasis of breast cancer from two patients and of prostate cancer from nine patients. (A) UMAP with scVelo streamlines. (B) Automatic cell type annotation using CellAssign. (C-F) Feature Plots of original cancer type (C), of mitochondrial percentage (D), of cell cycle (E), of copy number variant (CNV) score calculated using infercnvpy (F) and of normal or tumor information based on CNV score (G). (H) Dot plot of gene expression in the skeletal gene set. (I) Feature plots of gene expression related to osteoclast (OC) differentiation factors. (J, K) Dot plot (J) and Feature plots (K) of OC-related terms. (L) Cumulative bar plot of cell types using CellAssign. OC-enriched cluster 18 is highlighted. (M) Cumulative bar plot of cell type by patient. (N) Cumulative bar plot of normal or tumor by original cancer and by cell type.

**Figure S11.**
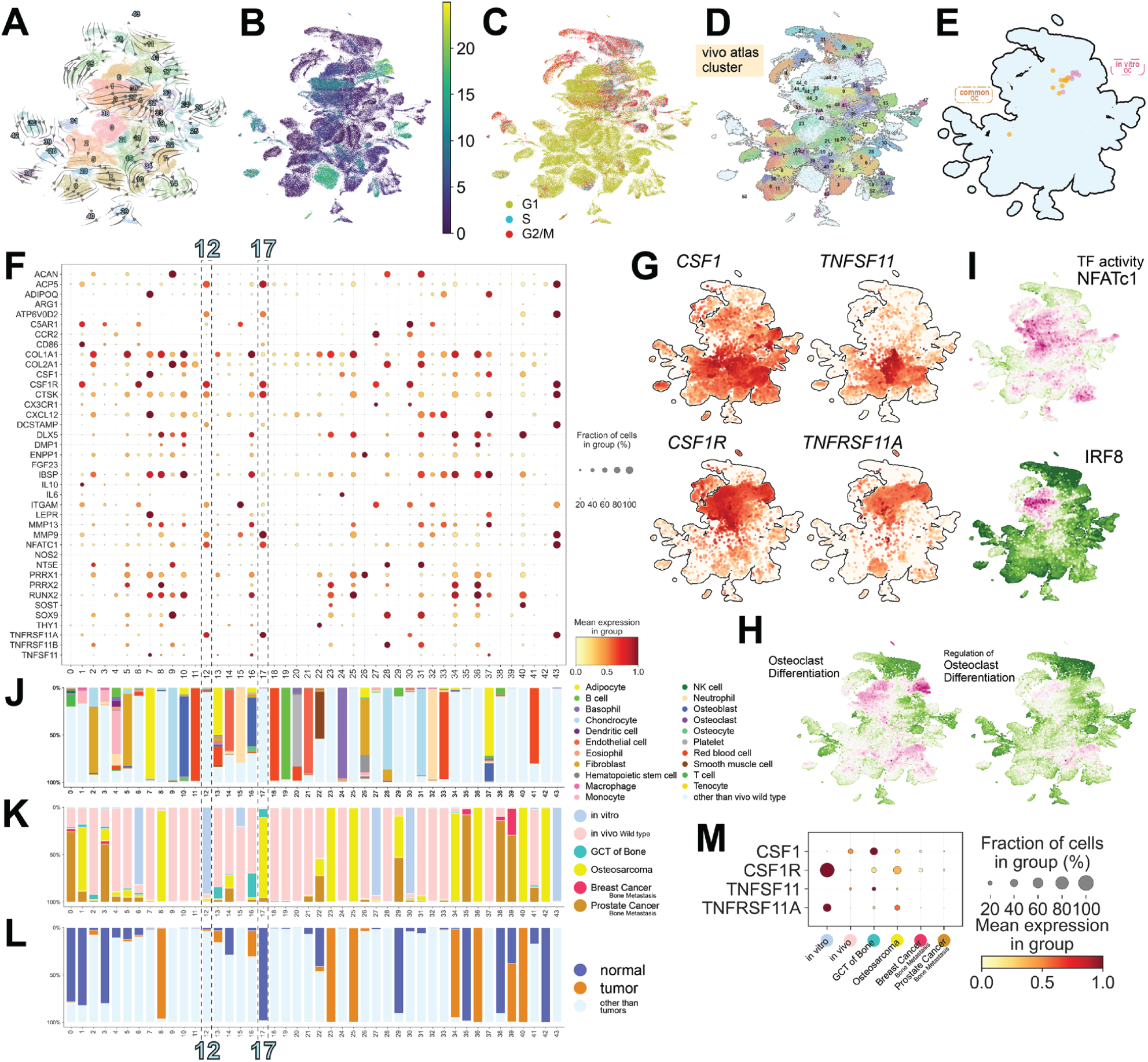
Basic parameters of final inclusive iSCseq with human bone tumors and metastasis (Figure 5). (A) UMAP of final inclusive iSCseq with humanized *in vitro*, humanized *in vivo* bone atlas and human bone tumor and metastasis datasets with scVelo streamlines. (B, C) Feature Plots of mitochondrial percentage (B) and of cell cycle (C). (D) *In vivo* bone atlas cells (Figure S4A) with cluster numbers. (E) *In vitro* osteoclast (OC) and common OC spots defined in Figure 4F in the enlarged view. (F) Dot plot of skeletal gene set. (G-I) Feature plots of gene expression related to OC differentiation factors (G), of gene ontology analysis (H) and of activities of transcriptional factors calculated using decoupleR and AUCell (I). (J-L) Cumulative bar plot of cell types by CellAssign (J), of sample categories (K) and of normal or malignant information in tumor samples (L). OC-enriched clusters 12 and 17 in final inclusive iSCseq are enclosed by dotted rectangles. (M) Dot plot of OC differentiation factors and their receptors according to sample category.

**Figure S12.**
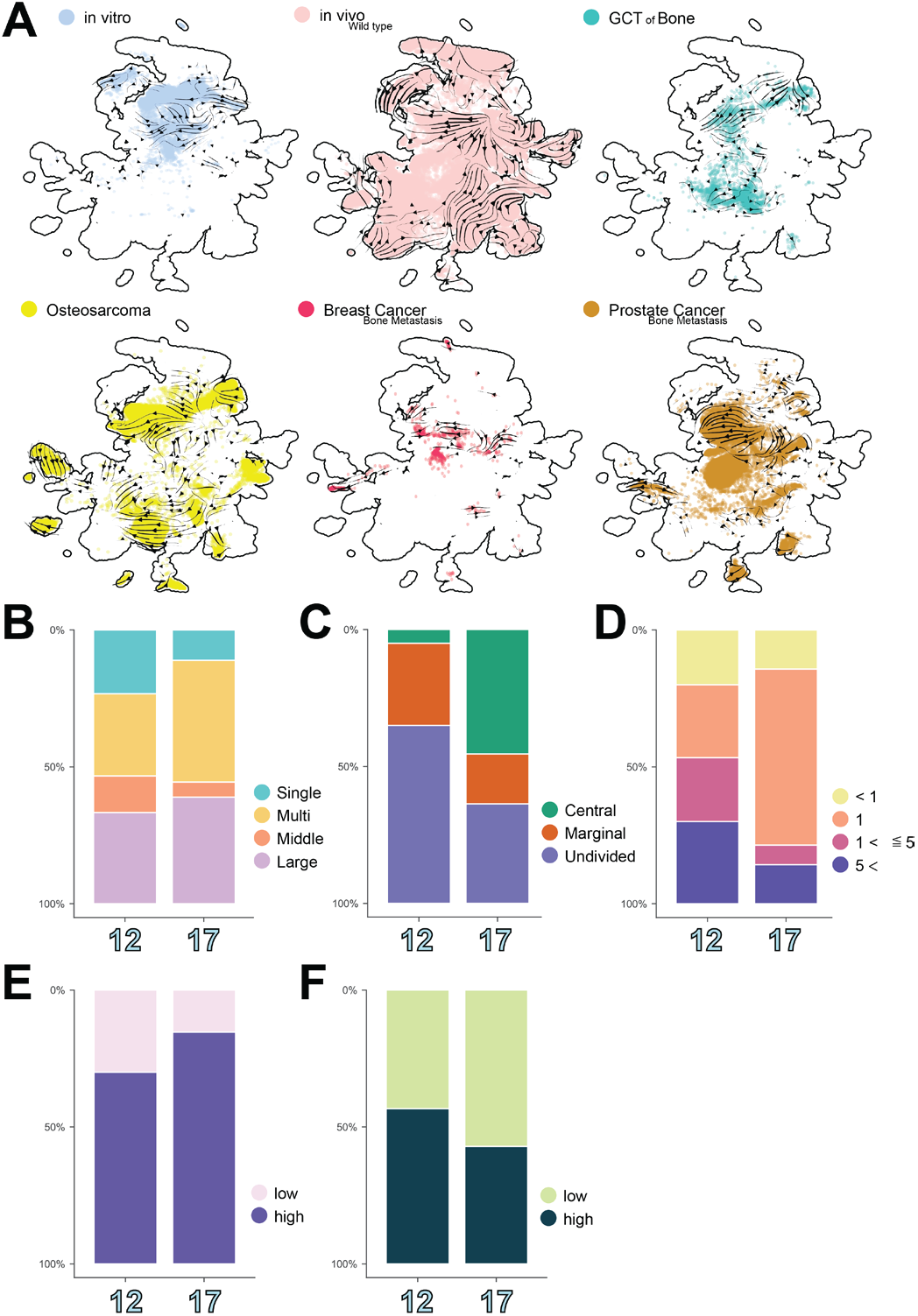
Basic characteristics of final inclusive iSCseq (Figure 5). (A) Cells in each sample category with scVelo streamlines. (B-F) Cumulative bar plots of the basic characteristics discussed in Figure 2, 3 and S3 between osteoclast-enriched clusters 12 and 17. (B) Nuclear Rank: bin, the number of nuclei in the origin of the cell. Large, 20 or more nuclei; Middle, 5 to 19 nuclei; Multi, 2 to 4 nuclei; Single, 1. (C) Positional information: Central or Marginal or Undivided in OC. (D) The binned number of nuclei in each sample. (E, F) High or low intensity of mitochondria (E) and calcium (F) based on the intensity from live cell images.

**Figure S13.**
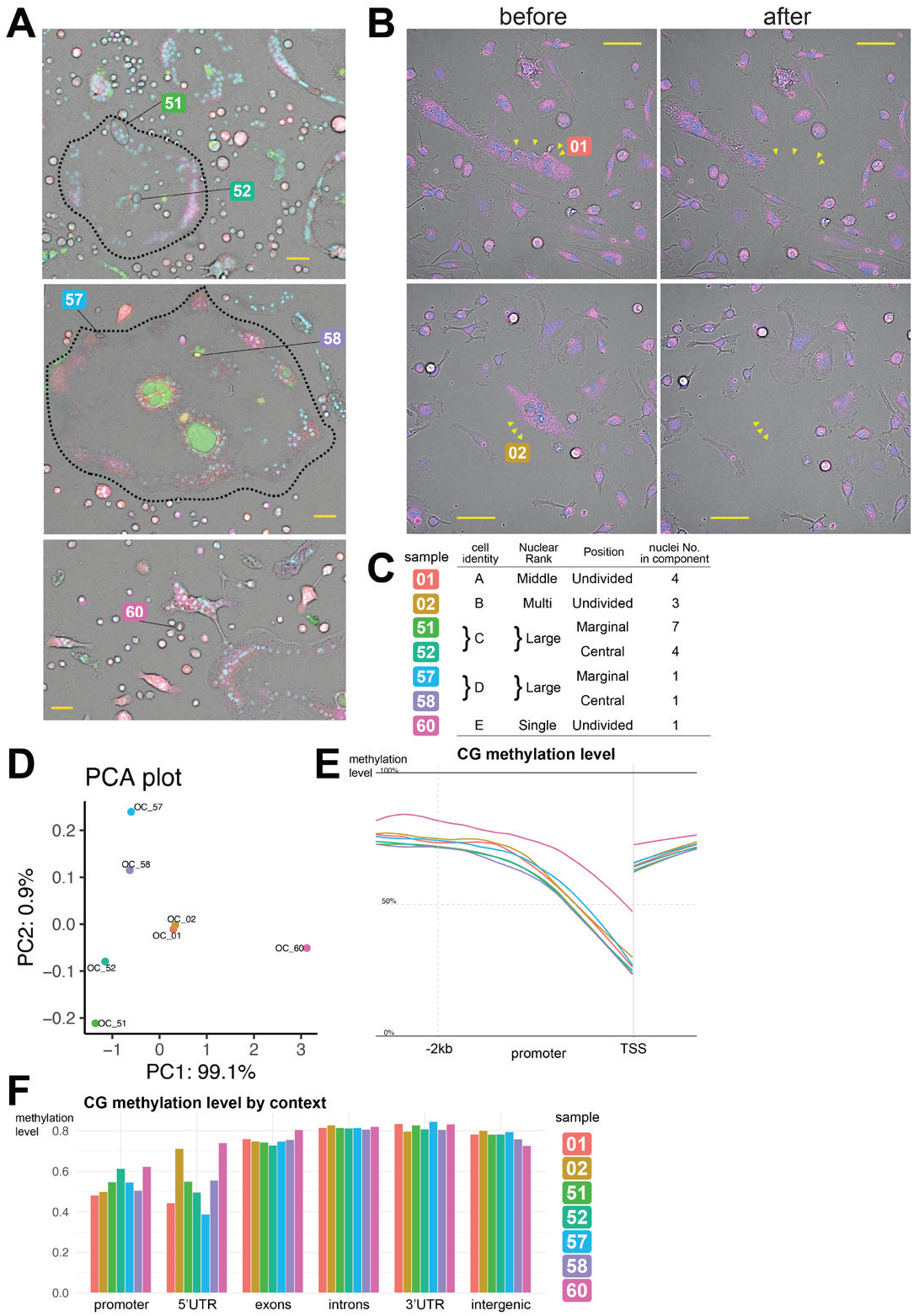
DNA methylation of sampled nuclei by whole genome bisulfite sequencing. (A) Geometrical information of targeted nuclei in large osteoclasts (O or a single-nucleated cell for whole genome bisulfite sequencing (WGBS). The OCs in the upper images are the same as those in Figure 1I. Blue: nuclei stained with Hoechst 33342; Green: Calcium stained with Fluo-4 NW Calcium Assay Kit; Red: acidic environment stained with pHrodo™ Red AM Intracellular pH Indicator; Purple: Tubulin stained with SiR-tubulin kit. Bar = 50 μm. (B) Sampling cases of nuclei for WGBS om multi-nucleated cells with Nuclear Rank middle (upper) or multi (lower), as defined in Figure 1J and 1K. Blue: Nuclei stained with Hoechst 33342; Red: CalbryteTM 590 ABD. Bar = 50 μm. (C) Basic operties of the sampled nuclei for WGBS. (D) Principal component analysis of DNA methylation levels n alized to library size. (E) Metagene plot of cytosine methylation levels in the CG context focus on the promoter region. TSS: Transcription Start Site. (F) Cytosine methylation levels in the CG context based on geno annotation.

**Figure S14.**
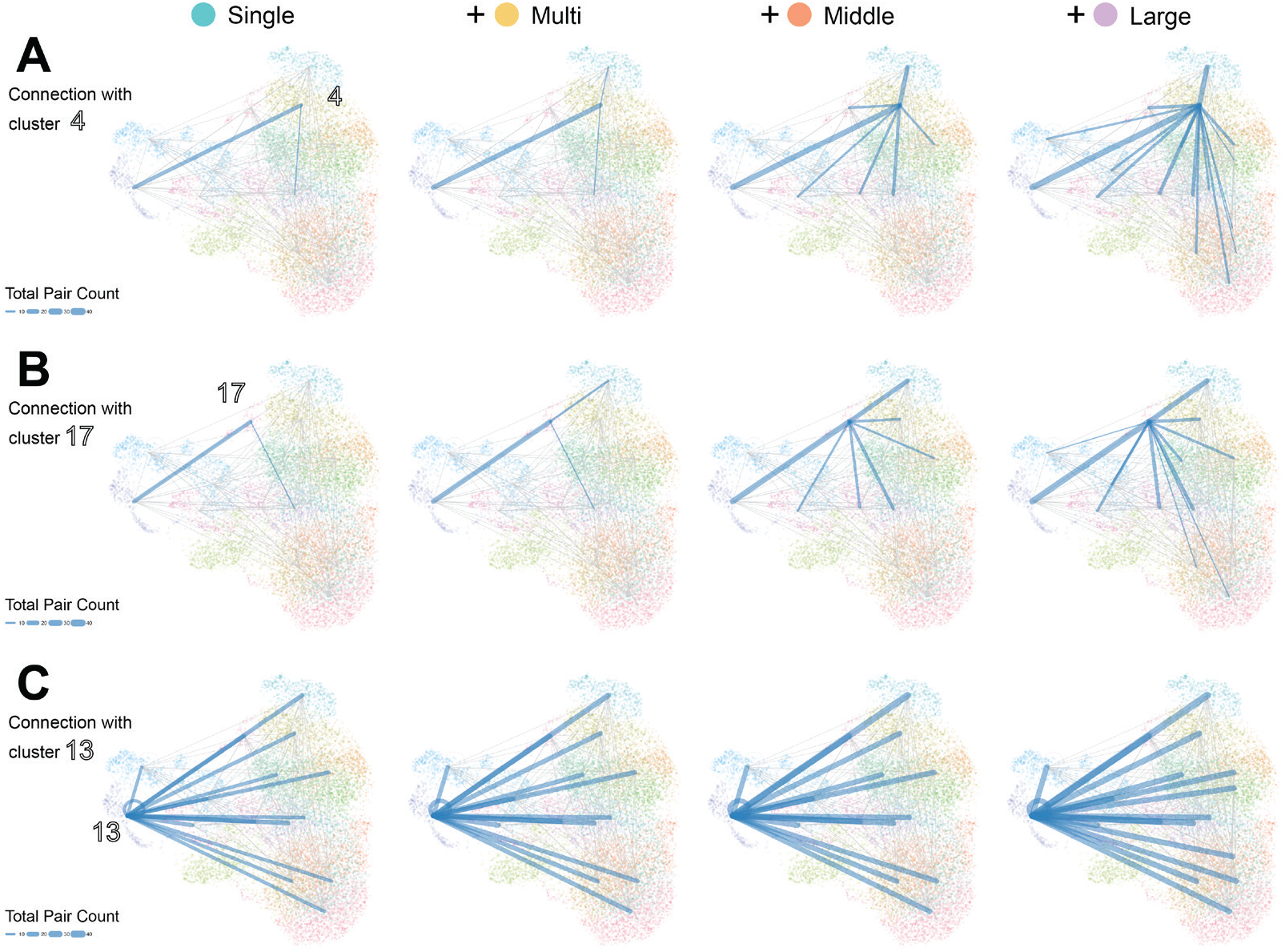
Colocalization of iSCseq components on *in vitro* iSCseq UMAP (Figure 6). Linkage of iSCseq components on inclusive *in vitro* iSCseq UMAP (Figure 2A) by gradually increasing Nuclear Rank. Edges and self-loops connected to osteoclast-enriched clusters 4 and 17 (A, B) and neutrophil-enriched cluster 13 (C). Colocalization of iSCseq components between different clusters as a node and within the same cluster as a self-loop. The edge or loop linewidth reflects the total number of pairs (10 or more highlighted). Nuclear Rank: bin, the number of nuclei in the origin of the cell. Large, 20 or more nuclei; Middle, 5 to 19 nuclei; Multi, 2 to 4 nuclei; Single, 1.

**Figure S15.**
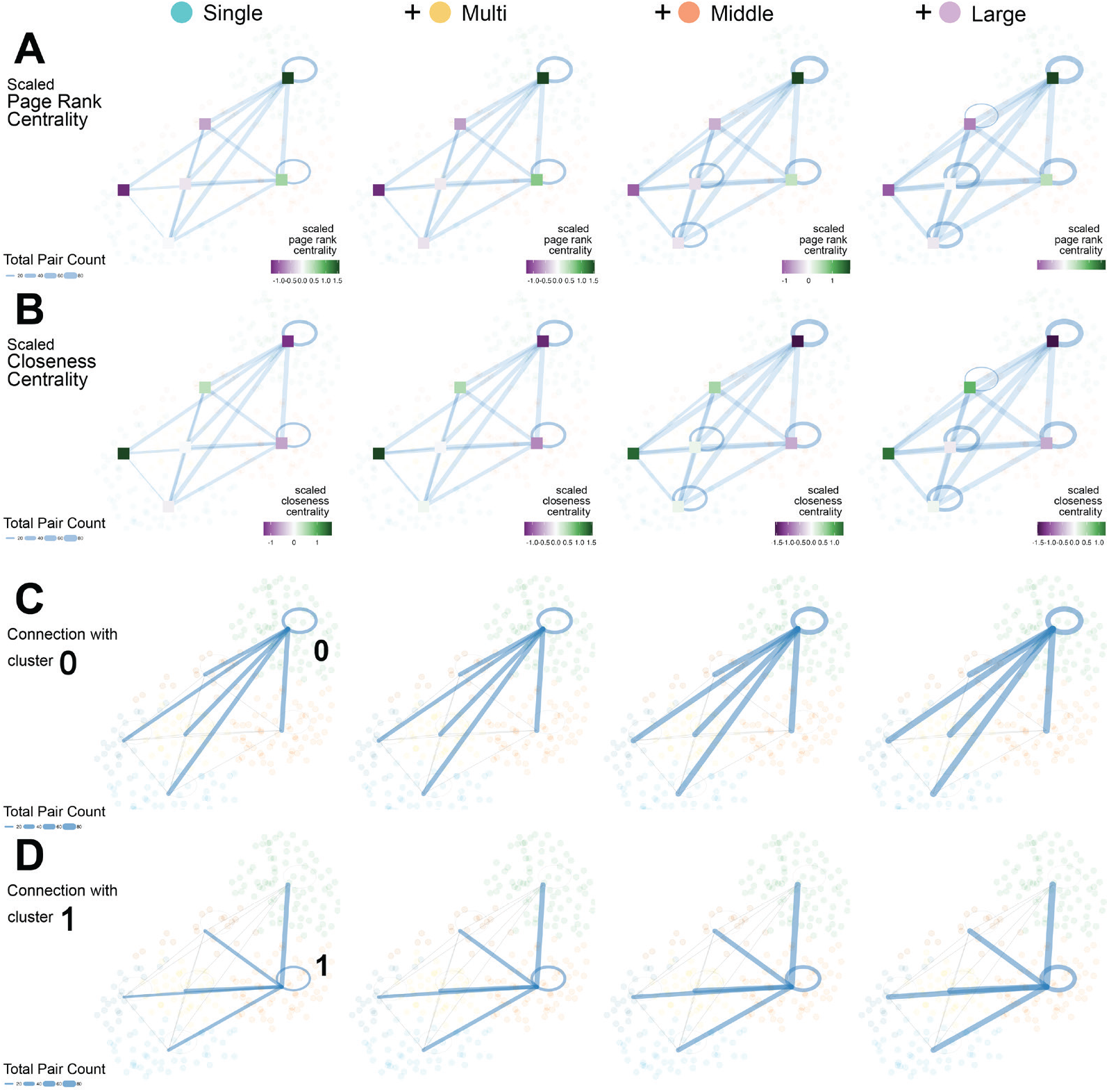
Colocalization of iSCseq components with centrality analysis on iSCseq alone UMAP. (A, B) Page rank (A) and closeness (B) centrality analysis with linkage of iSCseq alone components on iSCseq alone UMAP (Figure 1E) by gradually increasing Nuclear Rank. Colocalization of iSCseq components between different clusters as a node and within the same cluster as a self-loop. The edge or loop linewidth reflects the total number of pairs (20 or more highlighted). Nuclear Rank: bin, the number of nuclei in the origin of the cell. Large, 20 or more nuclei; Middle, 5 to 19 nuclei; Multi, 2 to 4 nuclei; Single, 1. (C, D) Edges and self-loops connected to osteoclast-enriched clusters 0 and 1.

## STAR ⋆ METHODS

### KEY RESOURCES TABLE

| REAGENT or RESOURCE | SOURCE | IDENTIFIER |
| --- | --- | --- |
| Antibodies |  |  |
| N/A |  |  |
| Bacterial and virus strains |  |  |
| N/A |  |  |
| Biological samples |  |  |
| N/A |  |  |
| Chemicals, peptides, and recombinant proteins |  |  |
| M-CSF, Murine, Recombinant, Animal Free | PeproTech | AF-315-02-10 µg |
| RANK Ligand soluble(sRANKL), Mouse, recombinant | Fujifilm Wako Pure Chemical corporation | 184-01791 10 µg |
| GM-CSF, Mouse, Recombinant | R&D Systems | 415-ML-010 10 µg |
| IL-4, Mouse, Recombinant | R&D Systems | 404-ML-010 10 µg |
| SiR-tubulin Kit | Cytoskeleton | CY-SC503 |
| Hoechst 33342 | Thermo Fisher | H3570 |
| MitoBright LT Deep Red | Dojindo | MT12 |
| Fluo-4 NW Calcium Assay Kit | Invitrogen | F36206 |
| pHrodo™ AM Ester | Thermo Fisher | P35372 |
| SiR-tubulin Kit | Cytoskeleton | CY-SC002 |
| DNA digestion buffer | Zymo Research | E1010-1-4 |
| AMPure XP PCR purification kit | Beckman Coulter | A63880 |
| Calbryte™ 590AM | AAT Bioquest | 20701 |
| Critical commercial assays |  |  |
| Single Cellome™ System SS2000 | Yokogawa Electric Corporation | SS2000 |
| SMART-Seq Single Cell PLUS kit | Takara Bio | R400751 |
| Bioanalyzer High Sensitivity DNA kit | Agilent | 5067-4626 |
| Pico Methyl-Seq Library Prep Kit | Zymo Research | D5456 |
| Deposited data |  |  |
| scRNA-seq of *in vitro* osteoclast differentiation post RANKL day0, 1, 3 | Tsukasaki *et al.*, 2020 | GSM4419695, 699 and 703 |
| scRNA-seq of *in vitro* osteoclast differentiation, wild type, M-CSF or M-CSF + RANKL | Canalis *et al.*, 2023 | GSM7906305 and GSM7906307 |
| *In vivo* bone scRNA-seq, reference 30 | Baryawno *et al.*, 2019 | GSM3674224-9 |
| *In vivo* bone scRNA-seq, reference 35 | Tikhonova *et al.*, 2019 | GSM2915578 |
| *In vivo* bone scRNA-seq, reference 29 | Baccin *et al.*, 2020. | GSM3466897 |
| *In vivo* bone scRNA-seq, reference 37 | Zhong *et al.*, 2020 | GSM4318799-802 |
| *In vivo* bone scRNA-seq, reference 36 | Wang *et al.*, 2021 | GSM4677821 |
| *In vivo* bone scRNA-seq, reference 33 | Matsushita *et al.*, 2020 | GSM4064321 and GSM4064323 |
| *In vivo* bone scRNA-seq, reference 32 | Johnston *et al.*, 2020 | GSM 4314099-101 |
| *In vivo* bone scRNA-seq, reference 34 | Mei *et al.*, 2022 | GSM6257588 |
| *In vivo* bone scRNA-seq, reference 31 | Hanai *et al.*, 2023 | GSM6820990 |
| *In vivo* bone scRNA-seq, reference 38 | Sivaraj et al., 2022 | GSM4667551-4 |
| giant cell tumor of bone scRNA-seq | Khazaei et al., 2020 | GSM4492128-31 |
| osteosarcoma scRNA-seq | Liu et al., 2021 | GSM4952363-5 and GSM5155198-200 |
| bone metastasis of breast cancer scRNA-seq | Ding et al., 2022 | GSM6870693-4 |
| bone metastasis of prostate cancer scRNA-seq | Kfoury et al., 2021 | GSM4274678, 82-3, 86, 89, 92, 97, 98 and 701 |
| Experimental models: Cell lines |  |  |
| N/A |  |  |
| Experimental models: Organisms/strains |  |  |
| Mouse/ C57BL/6J, wild type, female, 10-12 w.o. | Oriental Kobo Industrial Company | N/A |
| Oligonucleotides |  |  |
| N/A |  |  |
| Recombinant DNA |  |  |
| N/A |  |  |
| Software and algorithms |  |  |
| fastp ver. 0.23.2 | Chen et al., 2018 |  |
| STAR ver. 2.7.10b | Dobin et al., 2013 |  |
| STARsolo | Kaminow et al., 2021 |  |
| RSEM ver. 1.3.3 | Li and Dewey, 2011 |  |
| Scripts for NGS analysis by Nakato, R. | Nakato, 2023 |  |
| Python 3.10.11 | Van Rossum et al., 2009 |  |
| Scanpy ver. 1.9.4 | Wolf et al., 2018 |  |
| scverse | Virshup et al., 2023 |  |
| Seurat ver. 5.0.1 | Hao et al., 2021, 2022 |  |
| Scvi-tools ver. 1.0.2 | Gayoso et al., 2022 |  |
| Uniform Manifold Approximation and Projection | Leland McInnes, 2020 |  |
| scvelo ver. 0.2.5 | Bergen et al., 2020 |  |
| CellAssign | Zhang et al., 2019 |  |
| CellMarker 2.0 | Hu et al., 2023 |  |
| PanglaoDB | Franzen et al., 2019 |  |
| decoupleR ver. 1.4.0 | Badia et al., 2022 |  |
| AUCell ver. 1.18.1 | Aibar et al., 2017 |  |
| Mouse Gene Set Resources, Mouse MSigDB v2023.1 | Subramanian et al., 2005 |  |
| BioMart | Smedley et al., 2009 |  |
| infercnv (infercnvpy ver. 0.4.2) | the Trinity CTAT Project, 2020 |  |
| R v 4.3.0 | R Core team, 2023 |  |
| ggplot2 ver. 3.4.4 | Wickham, 2009 |  |
| Fiji / ImageJ2 ver. 2.14.0/1.54f | Schindelin et al., 2012 |  |
| Scipy ver. 1.1.2 | Virtanen et al., 2020 |  |
| Bismark ver. 0.22.3 | Krueger and Andrews, 2011 |  |
| HISAT2 ver 2.2.1 | Kin et al., 2019 |  |
| DeepTools ver. 3.5.5 | Ramirez et al., 2016 |  |
| GenomicRanges ver. 1.54.1 | Lawrence et al., 2013 |  |
| igraph v 2.0.2 | Csardi et al., 2024 |  |

## RESOURCE AVAILABILITY

### Lead contact

Further information and requests for resources and reagents should be directed to and will be fulfilled by the lead contact, Hiroyuki Okada.

### Data availability

Raw fastq files of iSCseq were deposited in DDBJ (DNA Data Bank of Japan). The accession numbers are DRA015536 and DRA016857. Raw fastq files of whole genome bisulfite sequencing will be public (accession numbers will be shared before publication.)

### Supplemental resources

Movie S1. Sampling a single nucleus in the peripheral zone of a large osteoclast (Figure 1A).

Movie S2. Sampling a single nucleus from four nuclei aggregation inside a middle-sized osteoclast (Figure 1B).

### Ethical considerations

Animal experiments were reviewed and approved by the University of Tokyo Institutional Animal Care and Use Committee (Permit Number: Med-P17-037).

## EXPERIMENTAL MODEL

### Mice

Wild-type female 10-to-12-week-old C57BL/6 mice were purchased from Oriental Kobo Industrial Company (Tokyo, Japan). The mice were euthanized by cervical dislocation to obtain bone marrow cells.

### Primary bone marrow cell cultures and *in vitro* osteoclast formation assay

Bone marrow cells of the lower limbs of mice were harvested in MEMalfa with L-Glutamine and Phenol Red (Fujifilm Wako Pure Chemical Corporation, Osaka, Japan) containing 10% fetal bovine serum (FBS; JRH Biosciences, Kansas, U.S.), 25 ng/mL macrophage colony-stimulating factor (M-CSF; R&D Systems, Minneapolis, MN, U.S.), and 1% penicillin-streptomycin (Thermo Fisher Scientific, Waltham, Massachusetts, U.S.) for three days to produce BMMs. To induce Osteoclasts, the BMMs were cultured on a 35 mm Glass Bottom Dish (Collagen Coat) f14 mm (Matsunami Glass, Osaka, Japan) in the presence of 50 ng/ml M-CSF (PeproTech Ltd., New Jersey, U.S.) and 50 ng/ml soluble RANKL (Fujifilm Wako Pure Chemical Corporation). To induce foreign body giant cells, BMMs were cultured with 50 ng/ml granulocyte-macrophage colony-stimulating factor (GM-CSF; R&D Systems, Minneapolis, MN, U.S.) and 50 ng/ml IL-4 (R&D Systems).

More than three nuclei with larger sizes compared to other types of cells were counted as morphological osteoclasts and FBGCs, according to the induction protocol.

## METHOD DETAILS

### Live cell staining

One hour before sampling, all nuclei were stained with 0.2 mg/mL Hoechst 33342, Trihydrochloride, Trihydrate (Invitrogen, Waltham, Massachusetts, U.S.) in PBS and incubated at 37°C for 10 min. Cells in Figure 1C, 2I, 2L, 3I and 3J were stained for nuclei with Hoechst 33342, then exposed to a mitochondrial staining solution containing MitoBright LT Deep Red (Dojindo, Kumamoto, Japan) in 1 mL Fluo-4 AM Assay buffer (HEPES buffer), incubated at 37°C for 45 min, and washed with MEMalfa containing 10% fetal bovine serum.

Cells in Figure 1I were stained for nuclei by Hoechst 33342, then stained with pHrodo™ AM Ester (Thermo Fisher) 2μL and PowerLoad™ concentrate (Thermo Fisher) 20 mL in 2 mL MEMalfa per 35 mm dish incubated at room temperature for 30 min. The pH probe was thoroughly washed with phosphate-buffered saline (PBS). Finally, the cells were loaded with the calcium indicator Fluo-4 (Fluo-4 NW Calcium Assay Kit, Invitrogen) and 500 nM SiR-tubulin Kit (Cytoskeleton, Denver, Colorado, U.S.) according to the manufacturer’s instructions.

Cells in Figure S3A and S3B were stained for nuclei by Hoechst 33342, then exposed to a calcium staining solution containing 1μL Calbryte™ 590AM (AAT Bioquest, California, U.S.) in 1 mL Fluo-4 AM Assay buffer, incubated at 37°C for 45 min, and washed with MEMalfa containing 10% fetal bovine serum. All cells sampled were incubated with the same reagents, including M-CSF and RANKL, under environmental conditions for differentiation.

### Sampling of intracellular components

Cellular components, including a single nucleus, were sampled using the Single CellomeTM System SS2000 (Yokogawa Electric Corporation, Japan). Before sampling, fine multi-colored images were obtained using confocal microscopy to confine the sampling target, including a single nucleus. When sampling cellular components, SS2000 finely tuned the position of probes in the Z-axis and sucked any cellular components into a 10 μm glass probe with the set negative pressure. The exact control of the tip with SS2000 is the key to collecting subcellular components from living cells. After sampling cellular components, probes maintain a small positive pressure so as not to pull in additional components, and finally carry onto microtubes with prechilled dissolution buffer at the right depth.

The buffer for RNA sequencing was 11.5 mL lysis buffer with 0.4% volume of RNase inhibitor. Both reagents were included in SMART-Seq Single Cell PLUS kit (Takara Bio, San Jose, CA, U.S.). Immediately after sampling in lysis buffer, microtubes containing samples were centrifuged briefly at 2,000 rpm and frozen at -80°C.

### Library construction for intra-single cell RNA-seq with micro input

SMART-Seq Single Cell PLUS kit (Takara Bio) was used for cDNA library construction, according to the manufacturer’s instructions. Frozen samples dissolved in lysis buffer with 0.4% volume of Ranse inhibitor in microtubes were thawed, 1μL 3’ SMART-Seq CDS Primer IIA was added to each microtube, then within the same microtubes incubated at 72°C for three minutes promptly. cDNA was synthesized according to the manufacturer’s instructions. In the cDNA amplification step, 20 PCR cycles were used, which was the maximum number of cycles required for the minimum input of RNA. The amplified cDNA was purified using an AMPure XP PCR purification kit (Beckman Coulter, Brea, California, U.S.). cDNA quality was checked using a Bioanalyzer High Sensitivity DNA kit (Agilent, Santa Clara, CA, U.S.) before library preparation. Samples with more than 10 times the cDNA concentration compared with the negative control by smear analysis within the 200-9,000 bp range were included.

SMART-Seq Library Prep Kit, including SMART-Seq Single-Cell PLUS Kit, was used for library preparation. The library amplification and PCR cycles were determined according to the concentrations detected by the bioanalyzer. After library amplification, the samples were pooled and purified once using AMPure. The pooled amplified library was qualified using a Bioanalyzer High-Sensitivity DNA kit.

### Library construction from sampled nuclei for whole genome bisulfite sequencing (WGBS)

Pico Methyl-Seq Library Prep Kit (Zymo research) was used for making WGBS libraries basically as manufacturer’s instruction. Because of the low input of DNA, such as a single nucleus, in the clean-up process, a higher concentration of DNA (3:1 ratio of DNA-binding buffer) was adopted. In the amplification process, 11 PCR cycles were performed, as previous paper^56^. In the amplification process with the index primer, Dual-indexed primers (5μM) 2.0μL in Unique dual index kit (Takara bio) was used instead of 0.5 μL Index Primer in the default Zymo kit. cDNA input was decreased as total volume was 25 μL. cDNA quality was qualified with Bioanalyzer High Sensitivity DNA kit (Agilent). After purification with 1.8x AMPure XP beads (Beckman Coulter), primer dimers around 60 bp were observed. We added a size selection of the cDNA library with 1.0x AMPure XP beads.

### Next generation sequencing of RNA-seq libraries

Pooled libraries were sequenced at 150 bp paired ends using HiSeq X Ten (Illumina, San Diego, CA, U.S.) by Macrogen Japan (Tokyo, Japan) or using NovaSeq 6000 by LiSDaC (Life Science Data Research Center at the University of Tokyo) and the Kashiwa sequencing team of the Platform for Advanced Genome Science, and by NGS core facility at the Research Institute for Microbial Diseases of Osaka University.

## QUANTIFICATION and STATISTICAL ANALYSES

### Bioinformatical analysis of intra-single cell RNA-seq (iSCseq)

All bioinformatics analyses were performed as we described previously^57^. Raw fastq files were mapped using STAR ver. 2.7.10b^58^ onto the primary assembly mouse genome GRCm39 and basic gene annotation vM31, or the human genome hg38 and basic annotation v42 downloaded from GENCODE^59^. The gene expression levels of iSCseq were summarized in a count matrix using RSEM^60^. Transcripts Per Kilobase Million (TPM) were calculated for iSCseq RSEM results using ‘rsem-generate-data-matrix-modified’ in script_rnakato^61^ prior to integration with 10x chromium datasets.

Downstream analyses were performed mainly using Python 3.10.11^62^. Scanpy^20^ in scverse project^22^ was used to process and visualize the count matrix. Scvi-tools were used to filter datasets considering basic parameters (cutoff values shown in Figs of Basic Parameters) and to integrate datasets with a model considering the experimental batch, including the author information of the dataset, total count, probe diameter, and cell cycle score based on cell cycle genes defined by the Seurat team^23,27,28,63,64^, percentages of leads on mitochondrial genes, and ribosomal genes. Scvi-tools are highly recommended for dataset integration in benchmark study^65^. The model was trained using a graphical processing unit (NVIDIA T4) on Google Clouds and Colab. Uniform Manifold Approximation and Projection (UMAP) was performed for dimension reduction19. Clustering was performed using the Leiden algorithm^66^ at the following resolution: 1 (Figure 1 and S1), 1 (Figure 2, 3, S2 and S3), 2 (Figure 4), 2.5 (Figure S4), 0.8 (Figure 5 and S11), 1.5 (Figure S8), 1 (Figure S9), and 1 (Figure S10). In bone milieu atlas processing (Figure 4A), we extracted cluster 44 containing osteoclasts and performed an additional clustering at a resolution of 1 into four subclusters, and then subcluster information was shadowed on the prior UMAP.

Automatic cell-type annotation was performed using CellAssign^26^, referring to the marker genes on CellMarker 2.0^67^ and PanglaoDB^68^. We classified the cells into six cell types, as shown in Figure 1H and 2D, for the *in vitro* assay. For the *in vivo* assay, we classified 21 cell types, as shown in Figure 4B, 5B, 5E and 5H. Gene ontology scores and regulatory transcriptional factor activities were calculated using decoupleR^25^ and AUcell^69,70^ referring to mSigDB^71-73^.

Prior to integrating human tumor scRNA-seq datasets, we converted mm39 mouse gene symbols into corresponding hg38 human gene symbols using homologous information from BioMart^74^. A total of 14,587 genes with human and mouse homologues were analyzed in the inclusive analysis in Figure 5, S10 and S11. Copy number variant (CNV) score was calculated using infercnvpy, which is a Python version of infercnv^75^. Based on the CNV score, we dichotomized normal or tumor cells.

M-A plots in Figure 4I,J were based on differentially expressed genes or terms analysis using scvi-tools and drawn using ggplot2^76^ on R environment^77^.

### Public resource of scRNA-seq datasets

In the inclusive iSCseq analysis with *in vitro* cultured osteoclast scRNA-seq, we used four datasets GSM4419695 (post RANKL stimulation day 0), GSM4419699 (day 1), GSM4419703 (day 3) by Tsukasaki et al.^15^, and wild type under three days treatment of M-CSF or M-CSF + RANKL by Canalis et al^17^.

For building *in vivo* bone and bone marrow atlas, we curated 32 scRNA-seq datasets using 10x chromium platform from ten papers including wild-type or wild-type equivalent control cells; 31,079 cells of GSM3674224-9 by Baryawno et al.^30^, 3,648 cells of GSM2915578 by Tikhonova et al.^35^, 1,165 cells of GSM3466897 by Baccin et al.^29^, 9,426 cells of GSM4318799-802 by Zhong et al.^37^, 1,572 cells of GSM4677821 by Wang et al.^36^, 3,869 cells of GSM4064321 and GSM4064323 by Matsushita et al.^33^, 6,445 cells of GSM 4314099-101 by Johnston et al.^32^, 3,804 cells of GSM6257588 by Mei et al.^34^, 3,535 cells of GSM6820990 by Hanai et al.^31^ and 9,755 cells of GSM4667551-4 by Sivaraj et al^38^. We downloaded fastq files and processed them by STARsolo mapping to the same mm39 reference genome.

We used giant cell tumor of bone scRNA-seq datasets (GSM4492128-31) from 4 patients by Khazaei et al.^40^, and osteosarcoma scRNA-seq datasets (GSM4952363-5 and GSM5155198-200) from 6 patients by Liu et al.^41^ In addition, scRNA-seq datasets from bone metastasis of breast cancer (GSM6870693-4) from 2 patients by Ding et al.^42^ and of prostate cancer (GSM4274678, 82-3, 86, 89, 92, 97, 98 and 701) from nine patients by Kfoury et al.^43^ were integrated.

### Analysis of images

The obtained images were processed and reconstructed using Fiji^78^. Enhanced contrast was used to reduce background noise. High or low signal intensities were determined manually by comparison with other surrounding cells or components inside the same cell.

### Bioinformatical analysis of DNA methylation by whole genome bisulfite sequencing

Raw fastq files were preprocessed and trimmed using fastp ver. 0.23.2^79^. Bismark ver. 0.22.3 was used as DNA methylation caller^80^. Reference genome for DNA methylation call was constructed from the primary assembly mouse genome GRCm39, then aligned using Bismark internally with HISAT2^81^ ver. 2.2.1 setting non_directional option according to Zymo Pico-Methyl technique. DNA methylation was extracted by bismark_methylation_extractor function ignoring the first 10 bp from both reads and making cytosine_report.

All Bismark methylation reports were assembled and normalized by library size. General normalized cytosine methylation level was assessed with principal component analysis. Then, we focused on CG context considering calculation cost. Metagene plot around transcription start site was calculated using deeptools^82^ ver. 3.5.5 and LOESS local regression was performed. Genomic annotation was performed using GenomicRanges^83^ and its related packages referring to mouse genome GRCm39 and basic gene annotation vM31. All figures were visualized with ggplot2.

### Network and centrality analysis of iSCseq components

First, we examined co-localization of each iSCseq sample based on live cell image data. iSCseq sample data were then matched to *in vitro* iSCseq cluster numbers and tabulated for each pair of cluster numbers. The cases in which cell populations of the same cluster were combined were also counted. Second, a colocalization map was created on the UMAP with the center of gravity of each cluster as a node, the union between clusters as a node, and the union within the same cluster as a self-loop. Third, page rank centrality and closeness centrality were calculated for each cluster to extract the core cluster of connections using igraph^84^. All network analyses’ images in Figure 6, S14 and S15 were made using ggplot2.

### Statistics

One-way ANOVA and post-hoc Tukey tests were performed using scipy^85^ to detect significant differences among the groups. If the p-value was less than 0.01, it was considered significant.

## REFERENCES

1. Luecken, M.D., and Theis, F.J. (2019). Current best practices in single-cell RNA-seq analysis: a tutorial. Mol Syst Biol 15, e8746. 10.15252/msb.20188746.

2. Wu, S.Z., Al-Eryani, G., Roden, D.L., Junankar, S., Harvey, K., Andersson, A., Thennavan, A., Wang, C., Torpy, J.R., Bartonicek, N., et al. (2021). A single-cell and spatially resolved atlas of human breast cancers. Nat Genet 53, 1334–1347. 10.1038/s41588-021-00911-1.

3. Rao, A., Barkley, D., Franca, G.S., and Yanai, I. (2021). Exploring tissue architecture using spatial transcriptomics. Nature 596, 211–220. 10.1038/s41586-021-03634-9.

4. Chen, W., Guillaume-Gentil, O., Rainer, P.Y., Gäbelein, C.G., Saelens, W., Gardeux, V., Klaeger, A., Dainese, R., Zachara, M., Zambelli, T., et al. (2022). Live-seq enables temporal transcriptomic recording of single cells. Nature. 10.1038/s41586-022-05046-9.

5. Tsukasaki, M., and Takayanagi, H. (2019). Osteoimmunology: evolving concepts in bone-immune interactions in health and disease. Nat Rev Immunol 19, 626–642. 10.1038/s41577-019-0178-8.

6. Tanaka, S., Tanaka, Y., Ishiguro, N., Yamanaka, H., and Takeuchi, T. (2018). RANKL: A therapeutic target for bone destruction in rheumatoid arthritis. Modern rheumatology 28, 9–16. 10.1080/14397595.2017.1369491.

7. Moller, M., Wolf, O., Bergdahl, C., Mukka, S., Rydberg, E.M., Hailer, N.P., Ekelund, J., and Wennergren, D. (2022). The Swedish Fracture Register - ten years of experience and 600,000 fractures collected in a National Quality Register. BMC Musculoskelet Disord 23, 141. 10.1186/s12891-022-05062-w.

8. Ferrari, S., and Langdahl, B. (2023). Mechanisms underlying the long-term and withdrawal effects of denosumab therapy on bone. Nat Rev Rheumatol 19, 307–317. 10.1038/s41584-023-00935-3.

9. Yasuda, H., Shima, N., Nakagawa, N., Yamaguchi, K., Kinosaki, M., Mochizuki, S., Tomoyasu, A., Yano, K., Goto, M., Murakami, A., et al. (1998). Osteoclast differentiation factor is a ligand for osteoprotegerin/osteoclastogenesis-inhibitory factor and is identical to TRANCE/RANKL. Proc Natl Acad Sci U S A 95, 3597–3602. 10.1073/pnas.95.7.3597.

10. Takayanagi, H., Kim, S., Koga, T., Nishina, H., Isshiki, M., Yoshida, H., Saiura, A., Isobe, M., Yokochi, T., Inoue, J., et al. (2002). Induction and activation of the transcription factor NFATc1 (NFAT2) integrate RANKL signaling in terminal differentiation of osteoclasts. Dev Cell 3, 889–901. 10.1016/s1534-5807(02)00369-6.

11. Tanaka, S., Amling, M., Neff, L., Peyman, A., Uhlmann, E., Levy, J.B., and Baron, R. (1996). c-Cbl is downstream of c-Src in a signalling pathway necessary for bone resorption. Nature 383, 528–531. 10.1038/383528a0.

12. Okada, H., Kajiya, H., Omata, Y., Matsumoto, T., Sato, Y., Kobayashi, T., Nakamura, S., Kaneko, Y., Nakamura, S., Koyama, T., et al. (2019). CTLA4-Ig Directly Inhibits Osteoclastogenesis by Interfering With Intracellular Calcium Oscillations in Bone Marrow Macrophages. J Bone Miner Res 34, 1744–1752. 10.1002/jbmr.3754.

13. Okada, H., Okabe, K., and Tanaka, S. (2020). Finely-Tuned Calcium Oscillations in Osteoclast Differentiation and Bone Resorption. Int J Mol Sci 22. 10.3390/ijms22010180.

14. Okada, H., and Tanaka, S. (2022). Plasmalemmal interface for calcium signaling in osteoclast differentiation. Curr Opin Cell Biol 74, 55–61. 10.1016/j.ceb.2022.01.001.

15. Tsukasaki, M., Huynh, N.C., Okamoto, K., Muro, R., Terashima, A., Kurikawa, Y., Komatsu, N., Pluemsakunthai, W., Nitta, T., Abe, T., et al. (2020). Stepwise cell fate decision pathways during osteoclastogenesis at single-cell resolution. Nat Metab 2, 1382–1390. 10.1038/s42255-020-00318-y.

16. Omata, Y., Okada, H., Uebe, S., Izawa, N., Ekici, A.B., Sarter, K., Saito, T., Schett, G., Tanaka, S., and Zaiss, M.M. (2022). Interspecies Single-Cell RNA-Seq Analysis Reveals the Novel Trajectory of Osteoclast Differentiation and Therapeutic Targets. JBMR Plus 6, e10631. 10.1002/jbm4.10631.

17. Canalis, E., Schilling, L., Yu, J., and Denker, E. (2023). NOTCH2 promotes osteoclast maturation and metabolism and modulates the transcriptome profile during osteoclastogenesis. The Journal of biological chemistry 300, 105613. 10.1016/j.jbc.2023.105613.

18. ten Harkel, B., Schoenmaker, T., Picavet, D.I., Davison, N.L., de Vries, T.J., and Everts, V. (2015). The Foreign Body Giant Cell Cannot Resorb Bone, But Dissolves Hydroxyapatite Like Osteoclasts. PLoS One 10, e0139564. 10.1371/journal.pone.0139564.

19. Leland McInnes, J.H., James Melville, (2020). UMAP: Uniform Manifold Approximation and Projection for Dimension Reduction. arXiv. 10.48550/arXiv.1802.03426.

20. Wolf, F.A., Angerer, P., and Theis, F.J. (2018). SCANPY: large-scale single-cell gene expression data analysis. Genome Biol 19, 15. 10.1186/s13059-017-1382-0.

21. Gayoso, A., Lopez, R., Xing, G., Boyeau, P., Valiollah Pour Amiri, V., Hong, J., Wu, K., Jayasuriya, M., Mehlman, E., Langevin, M., et al. (2022). A Python library for probabilistic analysis of single-cell omics data. Nat Biotechnol 40, 163–166. 10.1038/s41587-021-01206-w.

22. Virshup, I., Bredikhin, D., Heumos, L., Palla, G., Sturm, G., Gayoso, A., Kats, I., Koutrouli, M., Scverse, C., Berger, B., et al. (2023). The scverse project provides a computational ecosystem for single-cell omics data analysis. Nat Biotechnol. 10.1038/s41587-023-01733-8.

23. Hao, Y., Hao, S., Andersen-Nissen, E., Mauck, W.M., Zheng, S., Butler, A., Lee, M.J., Wilk, A.J., Darby, C., Zagar, M., et al. (2020). Integrated analysis of multimodal single-cell data. bioRxiv, 2020.2010.2012.335331. 10.1101/2020.10.12.335331.

24. Bergen, V., Lange, M., Peidli, S., Wolf, F.A., and Theis, F.J. (2020). Generalizing RNA velocity to transient cell states through dynamical modeling. Nat Biotechnol 38, 1408–1414. 10.1038/s41587-020-0591-3.

25. Badia, I.M.P., Velez Santiago, J., Braunger, J., Geiss, C., Dimitrov, D., Muller-Dott, S., Taus, P., Dugourd, A., Holland, C.H., Ramirez Flores, R.O., and Saez-Rodriguez, J. (2022). decoupleR: ensemble of computational methods to infer biological activities from omics data. Bioinform Adv 2, vbac016. 10.1093/bioadv/vbac016.

26. Zhang, A.W., O’Flanagan, C., Chavez, E.A., Lim, J.L.P., Ceglia, N., McPherson, A., Wiens, M., Walters, P., Chan, T., Hewitson, B., et al. (2019). Probabilistic cell-type assignment of single-cell RNA-seq for tumor microenvironment profiling. Nat Methods 16, 1007–1015. 10.1038/s41592-019-0529-1.

27. Hao, Y., Hao, S., Andersen-Nissen, E., Mauck, W.M., 3rd, Zheng, S., Butler, A., Lee, M.J., Wilk, A.J., Darby, C., Zager, M., et al. (2021). Integrated analysis of multimodal single-cell data. Cell 184, 3573–3587 e3529. 10.1016/j.cell.2021.04.048.

28. Hao, Y., Stuart, T., Kowalski, M., Choudhary, S., Hoffman, P., Hartman, A., Srivastava, A., Molla, G., Madad, S., Fernandez-Granda, C., and Satija, R. (2022). Dictionary learning for integrative, multimodal, and scalable single-cell analysis. bioRxiv. 10.1101/2022.02.24.481684.

29. Baccin, C., Al-Sabah, J., Velten, L., Helbling, P.M., Grunschlager, F., Hernandez-Malmierca, P., Nombela-Arrieta, C., Steinmetz, L.M., Trumpp, A., and Haas, S. (2020). Combined single-cell and spatial transcriptomics reveal the molecular, cellular and spatial bone marrow niche organization. Nat Cell Biol 22, 38–48. 10.1038/s41556-019-0439-6.

30. Baryawno, N., Przybylski, D., Kowalczyk, M.S., Kfoury, Y., Severe, N., Gustafsson, K., Kokkaliaris, K.D., Mercier, F., Tabaka, M., Hofree, M., et al. (2019). A Cellular Taxonomy of the Bone Marrow Stroma in Homeostasis and Leukemia. Cell 177, 1915–1932 e1916. 10.1016/j.cell.2019.04.040.

31. Hanai, A., Kawabata, A., Nakajima, K., Masuda, K., Urakawa, I., Abe, M., Yamazaki, Y., and Fukumoto, S. (2023). Single-cell RNA sequencing identifies Fgf23-expressing osteocytes in response to 1,25-dihydroxyvitamin D(3) treatment. Front Physiol 14, 1102751. 10.3389/fphys.2023.1102751.

32. Johnston, G., Ramsey, H.E., Liu, Q., Wang, J., Stengel, K.R., Sampathi, S., Acharya, P., Arrate, M., Stubbs, M.C., Burn, T., et al. (2020). Nascent transcript and single-cell RNA-seq analysis defines the mechanism of action of the LSD1 inhibitor INCB059872 in myeloid leukemia. Gene 752, 144758. 10.1016/j.gene.2020.144758.

33. Matsushita, Y., Nagata, M., Kozloff, K.M., Welch, J.D., Mizuhashi, K., Tokavanich, N., Hallett, S.A., Link, D.C., Nagasawa, T., Ono, W., and Ono, N. (2020). A Wnt-mediated transformation of the bone marrow stromal cell identity orchestrates skeletal regeneration. Nat Commun 11, 332. 10.1038/s41467-019-14029-w.

34. Mei, Y., Ren, K., Liu, Y., Ma, A., Xia, Z., Han, X., Li, E., Tariq, H., Bao, H., Xie, X., et al. (2022). Bone marrow-confined IL-6 signaling mediates the progression of myelodysplastic syndromes to acute myeloid leukemia. J Clin Invest 132. 10.1172/JCI152673.

35. Tikhonova, A.N., Dolgalev, I., Hu, H., Sivaraj, K.K., Hoxha, E., Cuesta-Dominguez, A., Pinho, S., Akhmetzyanova, I., Gao, J., Witkowski, M., et al. (2019). The bone marrow microenvironment at single-cell resolution. Nature 569, 222–228. 10.1038/s41586-019-1104-8.

36. Wang, J.S., Kamath, T., Mazur, C.M., Mirzamohammadi, F., Rotter, D., Hojo, H., Castro, C.D., Tokavanich, N., Patel, R., Govea, N., et al. (2021). Control of osteocyte dendrite formation by Sp7 and its target gene osteocrin. Nature Communications 12. 10.1038/s41467-021-26571-7.

37. Zhong, L., Yao, L., Tower, R.J., Wei, Y., Miao, Z., Park, J., Shrestha, R., Wang, L., Yu, W., Holdreith, N., et al. (2020). Single cell transcriptomics identifies a unique adipose lineage cell population that regulates bone marrow environment. Elife 9. 10.7554/eLife.54695.

38. Sivaraj, K.K., Majev, P.G., Jeong, H.W., Dharmalingam, B., Zeuschner, D., Schroder, S., Bixel, M.G., Timmen, M., Stange, R., and Adams, R.H. (2022). Mesenchymal stromal cell-derived septoclasts resorb cartilage during developmental ossification and fracture healing. Nat Commun 13, 571. 10.1038/s41467-022-28142-w.

39. La Manno, G., Soldatov, R., Zeisel, A., Braun, E., Hochgerner, H., Petukhov, V., Lidschreiber, K., Kastriti, M.E., Lonnerberg, P., Furlan, A., et al. (2018). RNA velocity of single cells. Nature 560, 494–498. 10.1038/s41586-018-0414-6.

40. Khazaei, S., De Jay, N., Deshmukh, S., Hendrikse, L.D., Jawhar, W., Chen, C.C.L., Mikael, L.G., Faury, D., Marchione, D.M., Lanoix, J., et al. (2020). H3.3 G34W Promotes Growth and Impedes Differentiation of Osteoblast-Like Mesenchymal Progenitors in Giant Cell Tumor of Bone. Cancer Discov 10, 1968–1987. 10.1158/2159-8290.CD-20-0461.

41. Liu, Y., Feng, W., Dai, Y., Bao, M., Yuan, Z., He, M., Qin, Z., Liao, S., He, J., Huang, Q., et al. (2021). Single-Cell Transcriptomics Reveals the Complexity of the Tumor Microenvironment of Treatment-Naive Osteosarcoma. Front Oncol 11, 709210. 10.3389/fonc.2021.709210.

42. Ding, K., Chen, F., Priedigkeit, N., Brown, D.D., Weiss, K., Watters, R., Levine, K.M., Heim, T., Li, W., Hooda, J., et al. (2022). Single cell heterogeneity and evolution of breast cancer bone metastasis and organoids reveals therapeutic targets for precision medicine. Ann Oncol 33, 1085–1088. 10.1016/j.annonc.2022.06.005.

43. Kfoury, Y., Baryawno, N., Severe, N., Mei, S., Gustafsson, K., Hirz, T., Brouse, T., Scadden, E.W., Igolkina, A.A., Kokkaliaris, K., et al. (2021). Human prostate cancer bone metastases have an actionable immunosuppressive microenvironment. Cancer Cell 39, 1464–1478 e1468. 10.1016/j.ccell.2021.09.005.

44. Yasui, T., Hirose, J., Tsutsumi, S., Nakamura, K., Aburatani, H., and Tanaka, S. (2011). Epigenetic regulation of osteoclast differentiation: possible involvement of Jmjd3 in the histone demethylation of Nfatc1. J Bone Miner Res 26, 2665–2671. 10.1002/jbmr.464.

45. Izawa, N., Kurotaki, D., Nomura, S., Fujita, T., Omata, Y., Yasui, T., Hirose, J., Matsumoto, T., Saito, T., Kadono, Y., et al. (2019). Cooperation of PU.1 With IRF8 and NFATc1 Defines Chromatin Landscapes During RANKL-Induced Osteoclastogenesis. J Bone Miner Res 34, 1143–1154. 10.1002/jbmr.3689.

46. Zhang, X., Li, T., Liu, F., Chen, Y., Yao, J., Li, Z., Huang, Y., and Wang, J. (2019). Comparative Analysis of Droplet-Based Ultra-High-Throughput Single-Cell RNA-Seq Systems. Mol Cell 73, 130–142 e135. 10.1016/j.molcel.2018.10.020.

47. Yanhong, L. (1998). Toward a qualitative search engine. IEEE Internet Computing 2, 24–29. 10.1109/4236.707687.

48. Brin, S., and Page, L. (1998). The anatomy of a large-scale hypertextual Web search engine. Computer Networks and ISDN Systems 30, 107–117. 10.1016/s0169-7552(98)00110-x.

49. Bavelas, A. (1950). Communication Patterns in Task-Oriented Groups. The Journal of the Acoustical Society of America 22, 725–730. 10.1121/1.1906679.

50. Hasegawa, T., Kikuta, J., Sudo, T., Matsuura, Y., Matsui, T., Simmons, S., Ebina, K., Hirao, M., Okuzaki, D., Yoshida, Y., et al. (2019). Identification of a novel arthritis-associated osteoclast precursor macrophage regulated by FoxM1. Nat Immunol 20, 1631–1643. 10.1038/s41590-019-0526-7.

51. Koga, T., Inui, M., Inoue, K., Kim, S., Suematsu, A., Kobayashi, E., Iwata, T., Ohnishi, H., Matozaki, T., Kodama, T., et al. (2004). Costimulatory signals mediated by the ITAM motif cooperate with RANKL for bone homeostasis. Nature 428, 758–763. 10.1038/nature02444.

52. Negishi-Koga, T., Gober, H.J., Sumiya, E., Komatsu, N., Okamoto, K., Sawa, S., Suematsu, A., Suda, T., Sato, K., Takai, T., and Takayanagi, H. (2015). Immune complexes regulate bone metabolism through FcRgamma signalling. Nat Commun 6, 6637. 10.1038/ncomms7637.

53. Terashima, A., Okamoto, K., Nakashima, T., Akira, S., Ikuta, K., and Takayanagi, H. (2016). Sepsis-Induced Osteoblast Ablation Causes Immunodeficiency. Immunity 44, 1434–1443. 10.1016/j.immuni.2016.05.012.

54. Paloneva, J., Mandelin, J., Kiialainen, A., Bohling, T., Prudlo, J., Hakola, P., Haltia, M., Konttinen, Y.T., and Peltonen, L. (2003). DAP12/TREM2 deficiency results in impaired osteoclast differentiation and osteoporotic features. J Exp Med 198, 669–675. 10.1084/jem.20030027.

55. Humphrey, M.B., Daws, M.R., Spusta, S.C., Niemi, E.C., Torchia, J.A., Lanier, L.L., Seaman, W.E., and Nakamura, M.C. (2006). TREM2, a DAP12-associated receptor, regulates osteoclast differentiation and function. J Bone Miner Res 21, 237–245. 10.1359/JBMR.051016.

56. Gravina, S., Dong, X., Yu, B., and Vijg, J. (2016). Single-cell genome-wide bisulfite sequencing uncovers extensive heterogeneity in the mouse liver methylome. Genome Biol 17, 150. 10.1186/s13059-016-1011-3.

57. Okada, H., Chung, U.I., and Hojo, H. (2023). Practical Compass of Single-Cell RNA-Seq Analysis. Curr Osteoporos Rep. 10.1007/s11914-023-00840-4.

58. Dobin, A., Davis, C.A., Schlesinger, F., Drenkow, J., Zaleski, C., Jha, S., Batut, P., Chaisson, M., and Gingeras, T.R. (2013). STAR: ultrafast universal RNA-seq aligner. Bioinformatics 29, 15–21. 10.1093/bioinformatics/bts635.

59. Frankish, A., Diekhans, M., Jungreis, I., Lagarde, J., Loveland, J.E., Mudge, J.M., Sisu, C., Wright, J.C., Armstrong, J., Barnes, I., et al. (2021). Gencode 2021. Nucleic Acids Res 49, D916–D923. 10.1093/nar/gkaa1087.

60. Li, B., and Dewey, C.N. (2011). RSEM: accurate transcript quantification from RNA-Seq data with or without a reference genome. BMC Bioinformatics 12, 323. 10.1186/1471-2105-12-323.

61. Nakato, R. (2023). Scripts for NGS analysis. https://github.com/rnakato/ script_rnakato.

62. Van Rossum, G., and Drake, F.L. (2009). Python 3 Reference Manual (CreateSpace).

63. Hafemeister, C., and Satija, R. (2019). Normalization and variance stabilization of single-cell RNA-seq data using regularized negative binomial regression. Genome Biol 20, 296. 10.1186/s13059-019-1874-1.

64. Choudhary, S., and Satija, R. (2022). Comparison and evaluation of statistical error models for scRNA-seq. Genome Biol 23, 27. 10.1186/s13059-021-02584-9.

65. Luecken, M.D., Buttner, M., Chaichoompu, K., Danese, A., Interlandi, M., Mueller, M.F., Strobl, D.C., Zappia, L., Dugas, M., Colome-Tatche, M., and Theis, F.J. (2022). Benchmarking atlas-level data integration in single-cell genomics. Nat Methods 19, 41–50. 10.1038/s41592-021-01336-8.

66. Traag, V.A., Waltman, L., and van Eck, N.J. (2019). From Louvain to Leiden: guaranteeing well-connected communities. Sci Rep 9, 5233. 10.1038/s41598-019-41695-z.

67. Hu, C., Li, T., Xu, Y., Zhang, X., Li, F., Bai, J., Chen, J., Jiang, W., Yang, K., Ou, Q., et al. (2023). CellMarker 2.0: an updated database of manually curated cell markers in human/mouse and web tools based on scRNA-seq data. Nucleic Acids Res 51, D870–D876. 10.1093/nar/gkac947.

68. Franzen, O., Gan, L.M., and Bjorkegren, J.L.M. (2019). PanglaoDB: a web server for exploration of mouse and human single-cell RNA sequencing data. Database (Oxford) 2019. 10.1093/database/baz046.

69. Aibar, S., and Aerts, S. (2016). AUCell: Analysis of ‘gene set’ activity in single-cell RNA-seq data. https://bioconductor.org/packages/release/bioc/html/AUCell. html.

70. Van de Sande, B., Flerin, C., Davie, K., De Waegeneer, M., Hulselmans, G., Aibar, S., Seurinck, R., Saelens, W., Cannoodt, R., Rouchon, Q., et al. (2020). A scalable SCENIC workflow for single-cell gene regulatory network analysis. Nat Protoc 15, 2247– 2276. 10.1038/s41596-020-0336-2.

71. Subramanian, A., Tamayo, P., Mootha, V.K., Mukherjee, S., Ebert, B.L., Gillette, M.A., Paulovich, A., Pomeroy, S.L., Golub, T.R., Lander, E.S., and Mesirov, J.P. (2005). Gene set enrichment analysis: a knowledge-based approach for interpreting genome-wide expression profiles. Proc Natl Acad Sci U S A 102, 15545–15550. 10.1073/pnas.0506580102.

72. Castanza, A.S., Recla, J.M., Eby, D., Thorvaldsdottir, H., Bult, C.J., and Mesirov, J.P. (2023). Extending support for mouse data in the Molecular Signatures Database (MSigDB). Nat Methods. 10.1038/s41592-023-02014-7.

73. Liberzon, A., Subramanian, A., Pinchback, R., Thorvaldsdottir, H., Tamayo, P., and Mesirov, J.P. (2011). Molecular signatures database (MSigDB) 3.0. Bioinformatics 27, 1739–1740. 10.1093/bioinformatics/btr260.

74. Smedley, D., Haider, S., Ballester, B., Holland, R., London, D., Thorisson, G., and Kasprzyk, A. (2009). BioMart--biological queries made easy. BMC Genomics 10, 22. 10.1186/1471-2164-10-22.

75. inferCNV of the Trinity CTAT Project (2020). inferCNV. https://github.com/broadinstitute/inferCNV.

76. Wickham, H. (2009). ggplot2: Elegant Graphics for Data Analysis (Springer-Verlag New York).

77. R Core Team (2023). R: A Language and Environment for Statistical Computing, .

78. Schindelin, J., Arganda-Carreras, I., Frise, E., Kaynig, V., Longair, M., Pietzsch, T., Preibisch, S., Rueden, C., Saalfeld, S., Schmid, B., et al. (2012). Fiji: an open-source platform for biological-image analysis. Nat Methods 9, 676–682. 10.1038/nmeth.2019.

79. Chen, S., Zhou, Y., Chen, Y., and Gu, J. (2018). fastp: an ultra-fast all-in-one FASTQ preprocessor. Bioinformatics 34, i884–i890. 10.1093/bioinformatics/bty560.

80. Krueger, F., and Andrews, S.R. (2011). Bismark: a flexible aligner and methylation caller for Bisulfite-Seq applications. Bioinformatics 27, 1571–1572. 10.1093/bioinformatics/btr167.

81. Kim, D., Paggi, J.M., Park, C., Bennett, C., and Salzberg, S.L. (2019). Graph-based genome alignment and genotyping with HISAT2 and HISAT-genotype. Nat Biotechnol 37, 907–915. 10.1038/s41587-019-0201-4.

82. Ramirez, F., Ryan, D.P., Gruning, B., Bhardwaj, V., Kilpert, F., Richter, A.S., Heyne, S., Dundar, F., and Manke, T. (2016). deepTools2: a next generation web server for deep-sequencing data analysis. Nucleic Acids Res 44, W160–165. 10.1093/nar/gkw257.

83. Lawrence, M., Huber, W., Pages, H., Aboyoun, P., Carlson, M., Gentleman, R., Morgan, M.T., and Carey, V.J. (2013). Software for computing and annotating genomic ranges. PLoS Comput Biol 9, e1003118. 10.1371/journal.pcbi.1003118.

84. Csardi, G., and Nepusz, T. (2006). The igraph software package for complex network research. InterJournal Complex Systems, 1695.

85. Virtanen, P., Gommers, R., Oliphant, T.E., Haberland, M., Reddy, T., Cournapeau, D., Burovski, E., Peterson, P., Weckesser, W., Bright, J., et al. (2020). SciPy 1.0: fundamental algorithms for scientific computing in Python. Nat Methods 17, 261–272. 10.1038/s41592-019-0686-2.

